# Biochemical Characterization of a Fungal Bifunctional Type I Diterpene Synthase Brings a Cryptic Natural Product to Light

**DOI:** 10.1101/2022.05.03.490360

**Authors:** Peng Zhang, Guangwei Wu, Stephanie C. Heard, Changshan Niu, Stephen A. Bell, Fengli Li, Ying Ye, Yonghui Zhang, Jaclyn M. Winter

## Abstract

We report the identification of the tnd biosynthetic cluster from the marine-derived fungal strain Aspergillus flavipes CNL-338 and in vivo characterization of a cryptic type I diterpene synthase. Heterologous expression of the bifunctional terpene synthase TndC in Saccharomyces cerevisiae led to the discovery of a new diterpene backbone, talarodiene, harboring a benzo[a]cyclopenta[d]cyclooctane tricyclic fused ring system. The cyclization mechanism that converts geranylgeranyl diphosphate to the tricyclic hydrocarbon skeleton was investigated using ^13^C-labeling studies and stable isotope tracer experiments showed the biotransformation of talarodiene into talaronoid C.

---

Terpenes are the largest class of natural products and are produced by all kingdoms of life. These compounds possess enormous structural diversity and exhibit various biological activities ranging from anticancer and antimalarial to carcinogens and mycotoxins.^1^ Despite their structural complexity, all terpenes are derived from the universal C_5_ hemiterpene precursors dimethylallyl diphosphate (DMAPP) and isopentenyl diphosphate (IPP). Coupling of these C_5_ precursors, facilitated by prenyltransferases (PTs), generates linear, achiral polyprenyl diphosphates that can be transformed by terpene cyclases (TCs) into complex scaffolds containing multiple fused rings and stereogenic centers.^2–6^ The structural diversity associated with terpenes often originates from the cyclization step, and TCs catalyze some of the most complex reactions in natural products chemistry.

In fungi, although condensation and cyclization reactions occur independently, bifunctional terpene synthases have been characterized where the C-terminal half is responsible for producing the polyprenyl diphosphate and the N-terminal half catalyzes the cyclization reaction. The first fungal type I diterpene synthase, PaFS, was characterized in 2007 from *Phomopsis amygdali* and shown to produce fusicoccin **1**.^7^ AcOS was the first type I sesterterpene synthase characterized in 2013 from *Aspergillus clavatus* and is responsible for the biosynthesis of ophiobolin F **2**.^8^ Because of their potential in synthesizing diverse hydrocarbon skeletons, subsequent genome mining efforts focused on identifying additional cryptic type I bifunctional terpene synthases. As a result, a number of fungal type I sesterterpene synthases were characterized.^9–17^ However, since the discovery of PaFS, only a limited number of type I diterpene synthases have been identified and include those responsible for the production of variediene **3**,^18^ phomopsene **4**,^19^ brassicicene **5**,^20^ a precursor to the cyclopiane-type diterpenes **6**, and dolasta-1 (15),8-diene **7** (Figure 1). Given our limited knowledge of type I diterpene synthases, the discovery and biochemical characterization of new enzymes would bring to light cryptic natural products, unveil novel cyclization reactions, and allow for more informed bioinformatic predictions. In this work, we describe the discovery and *in vivo* characterization of a cryptic bifunctional type I diterpene synthase from a marine-derived fungus that synthesizes a new tricyclic 5-8-6 hydrocarbon skeleton. The use of stable tracer isotope experiments also allowed us to show the biotransformation of the diterpene backbone into the talaronoid class of natural products.

**Figure 1.**
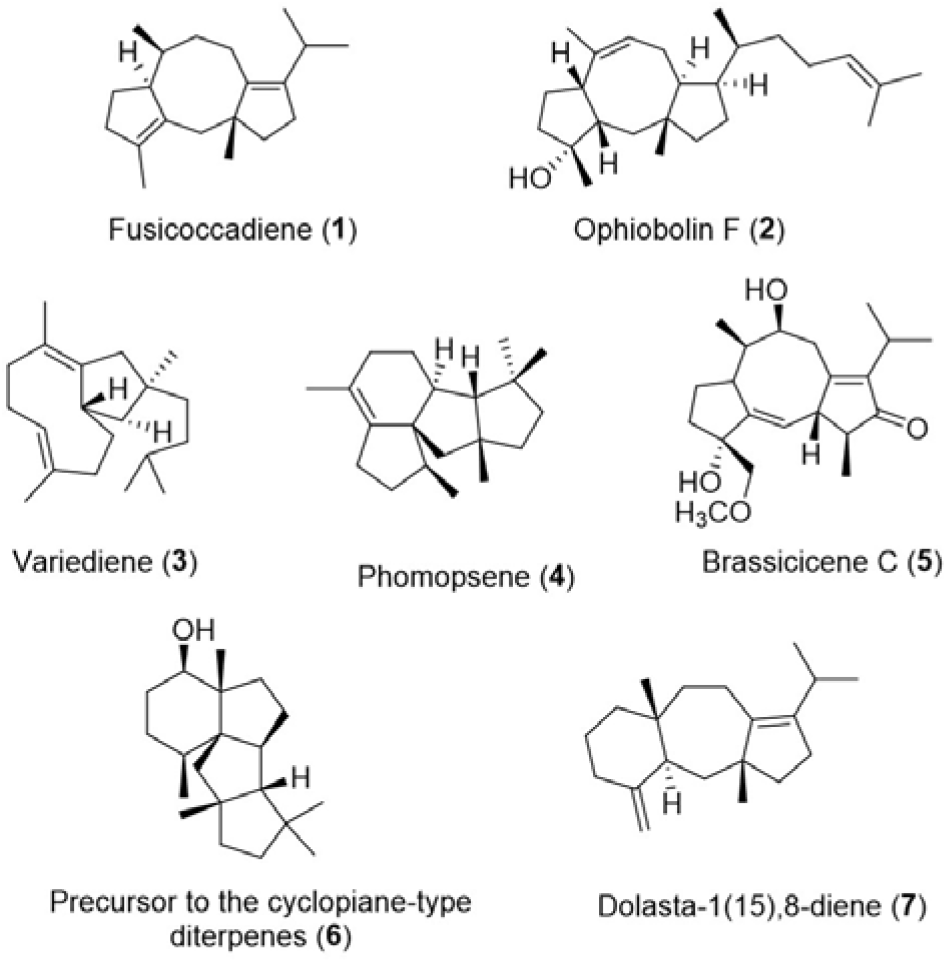
Structures of selected fungal diterpenes and sesterterpenes produced by type I bifunctional terpene synthases.

It is known that marine organisms are prolific producers of bioactive natural products and often produce molecules not observed in their terrestrial counterparts.^23^ The previously characterized type I bifunctional terpene synthases were identified exclusively from terrestrial fungi, and given the tremendous promise that marine organisms hold for characterizing novel biosynthetic enzymes, we turned to marine-derived fungi as an underexplored resource for identifying and characterizing type I terpene synthases. Recently, our group sequenced the genome of the marine-derived fungus *Aspergillus flavipes* CNL-338,^24^ and using the PaFS and AcOS sequences as probes, we scanned the genome for bifunctional terpene synthases. A 21 kb biosynthetic cluster harboring a cryptic chimeric synthase was identified (Figure 2a), and bioinformatic analysis of TndC revealed that the 764 amino acid protein possessed both PT and TC domains. Multiple sequence alignment also showed that TndC contains the conserved aspartate-rich DDxxD motif for Mg^2+^ binding in both the PT and TC domains, as well as a second NSE Mg^2+^-binding motif in the TC domain indicative of type I cyclases (Figure S2). Phylogenetic comparison of the cryptic chimeric synthase with known fungal-derived diterpene and sesterterpene synthases showed that TndC clades between PaFS and the sesterterpene synthase EvAS (Figure S1), suggesting that TndC could be producing a new terpene skeleton; however, it was not clear if the product was a diterpene or sesterterpene.

**Figure 2.**
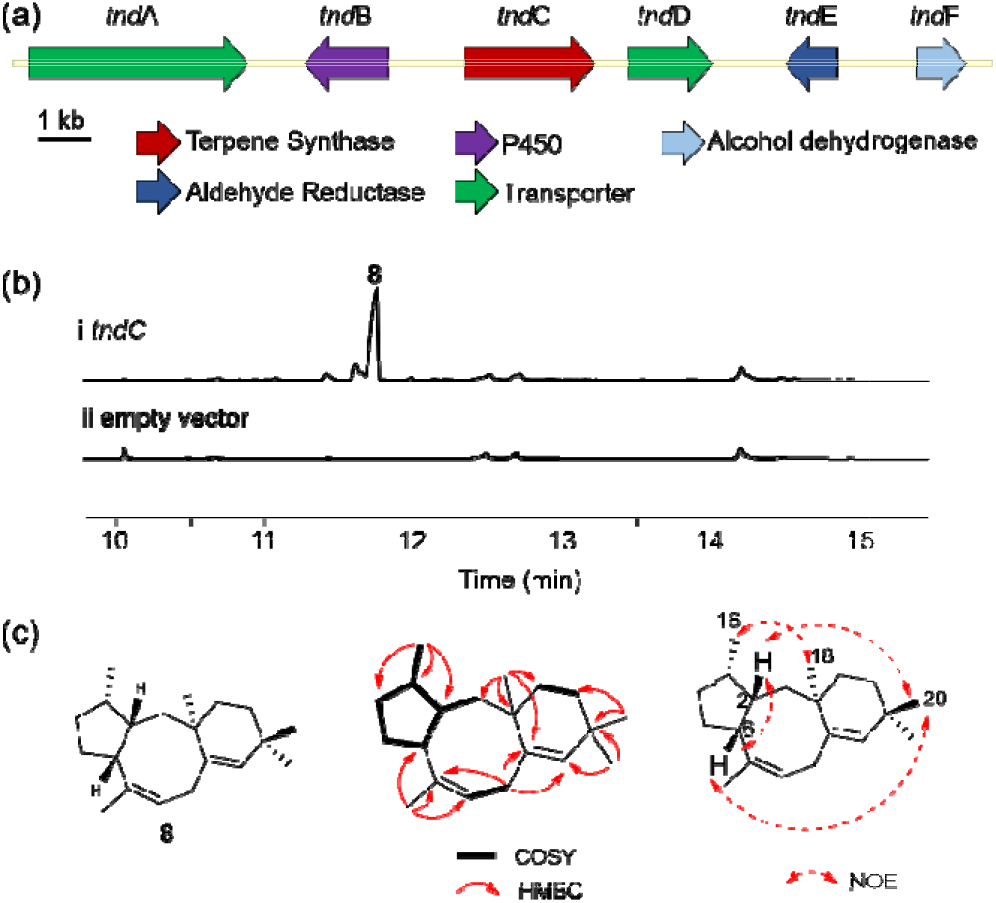
Characterization of the type I diterpene synthase *tndC* from *A. flavipes* CNL-338. (a) Organization of the *tnd* biosynthetic gene cluster in *A. flavipes* CNL-338. (b) GC-MS analysis (TIC) of extracts from *S. cerevisiae* ZXM144 transformed with (i) a plasmid-borne *tndC* or (ii) empty vector. (c) The structure identification of compound 8 and key 2D NMR correlations.

Initial efforts at expressing recombinant TndC from *Escherichia coli* and *Saccharomyces cerevisiae* failed to generate any soluble protein. Thus, to elucidate the product of TndC, we heterologously expressed intron-free *tnd*C in *Saccharomyces cerevisiae* ZXM144.^25^ GC-MS analysis of crude extracts of *S. cerevisiae* ZXM144 transformed with *tndC* compared to an empty vector control revealed the presence of a new product, **8**, with a m/z 272 [M]^+^ (Figures 2b and S7), supporting the production of a diterpene over a sesterterpene. HREIMS coupled 1D and 2D NMR spectroscopy (Figures S7 and S11-S15 and Table S2) identified that the planar structure of **8**, which was named talarodiene, contained a benzo[*a*]cyclopenta[*d*]cyclooctane tricyclic hydrocarbon backbone (Figure 2c). NOESY correlations were used to assign the relative configuration of **8** (Figures S16-S22), and ECD calculations (Figure S8) determined the absolute configuration to be (2S,3S,6R,11R).

With the isolation of **8**, the cyclization mechanism converting geranylgeranyl diphosphate (GGPP) into the 5-8-6 tricyclic hydrocarbon skeleton was investigated using ^13^C-labeling studies. [1-^13^C]acetate, [2-^13^C]acetate and [1,2-^13^C_2_]acetate were administered independently to *tnd*C-transformed *S. cerevisiae* ZXM144, and the corresponding labeling patterns of ^13^C-enriched **8** were analyzed by NMR (Figures S23-S25 and Table S3). From the [1,2-^13^C_2_]acetate labeling patterns, a cyclization mechanism is proposed in Scheme 1. Cleavage of diphosphate followed by 1,11-10,14-cyclization converts GGPP to the bicyclic tertiary cation intermediate **8a^+^**. Ring expansion of **8a^+^** from a 1,2 alkyl shift forms the cation intermediate **8b^+^**, which is transformed to the tertiary cation intermediate **8c^+^** following a transannular proton transfer. A 1,2-hydride shift and 2,6-cyclization forms intermediate **8d^+^**, and deprotonation at C-8 ultimately yields **8**.

**Scheme 1.**
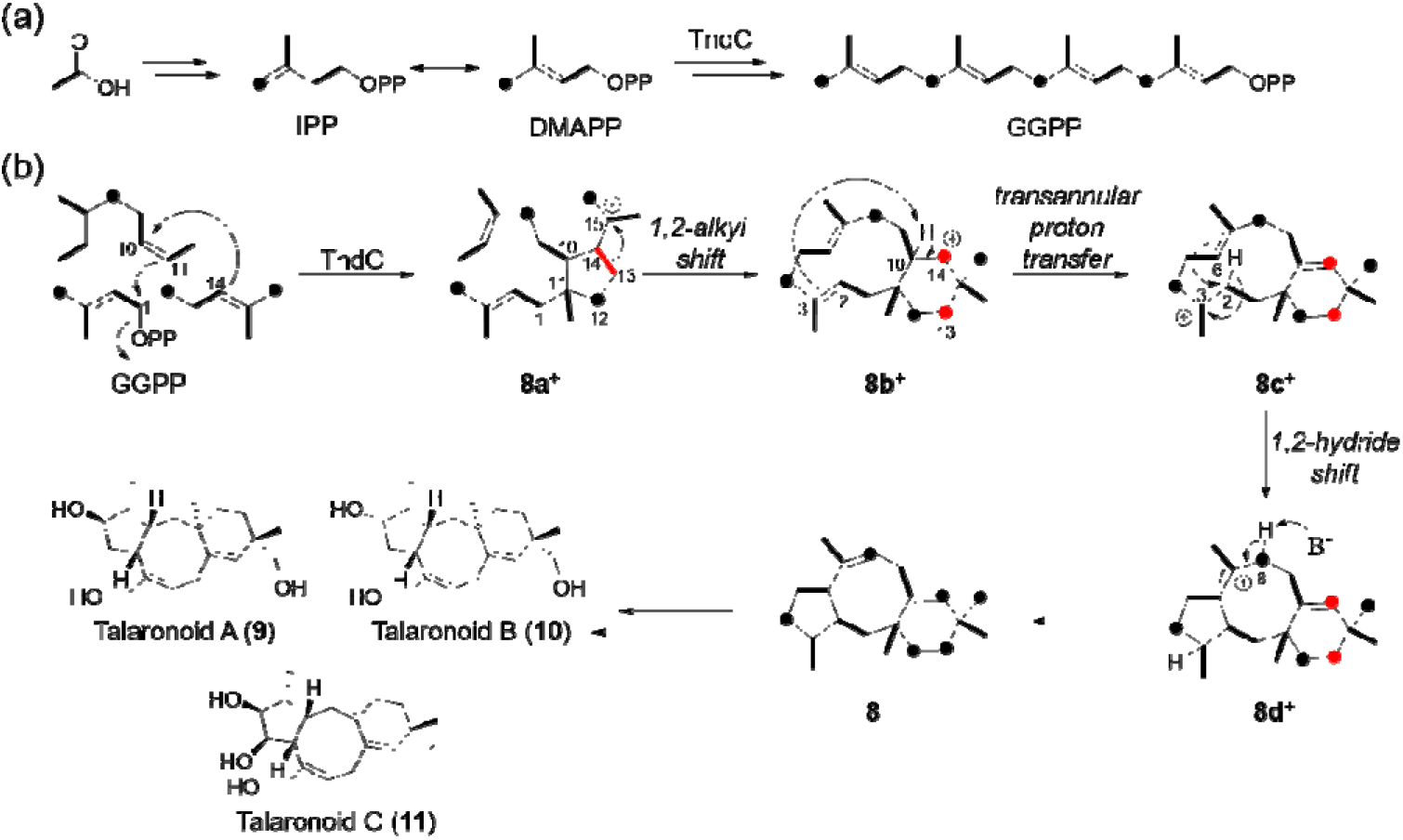
Proposed biosynthesis of the talarodiene backbone. (a) Biosynthesis of the linear achiral precursor geranylgeranyl diphosphate using the C-terminal prenyltransferase domain of TndC. (b) Formation of the 5-8-6 tricyclic talarodiene backbone **8** by the N-terminal cyclization domain of TndC. [1,2-13C2]acetate labeling patterns are shown as black bold lines and dots to signify double and single enrichments, respectively. Red dots indicate the C□C bond breakage of an intact acetate unit.

After heterologous expression of the cryptic *tnd*C gene led to the isolation of **8**, we turned back to the original host and evaluated *A. flavipes* CNL-338 for its production of this new tricyclic diterpene (Figure 3). Unfortunately, we were unable to detect the presence of **8** in crude extracts using GC-MS and LC-MS analyses, which suggests that 8 is an intermediate in the biosynthetic pathway and is modified by tailoring enzymes encoded in the *tnd* gene cluster. In addition to the diterpene synthase, the *tnd* cluster encodes several oxidative enzymes including the cytochrome P450 (*tnd*B), an aldehyde reductase (*tnd*E), and an alcohol dehydrogenase (*tnd*F) (Table S4). Given the type of tailoring enzymes present, we speculated that the cytochrome P450 TndB would be the next enzyme in the biosynthetic pathway. Indeed, GC-MS analysis of the Δ*tndB* mutant showed the accumulation of **8** (Figure 3 and S10).

**Figure 3.**
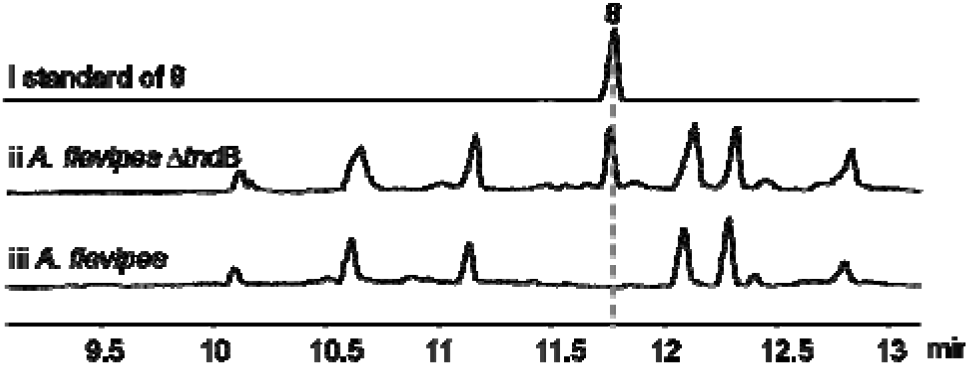
GC-MS chromatograms (TIC) of (i) a standard of compound **8**, (ii) crude extract of Δ*tndB* strain and (iii) crude extract from wild-type *A. flavipes* CNL-338.

While the gene inactivation experiments unequivocally linked the *tnd* biosynthetic cluster to **8** in *A. flavipes* CNL-338, the final natural product(s) produced by the pathway were unknown. Recently, a group of diterpenoids, talaronoid A **9**, B **10**, and C **11**, containing a 5-8-6 fused ring system were isolated from *Talaromyces stipitatus* (Scheme 1).^26^ Using the amino acid sequence of TndC, we identified a gene cluster in *T. stipitatus* harboring an assortment of genes similar to those in the *tnd* biosynthetic cluster (Figures S3-S5). However, when we scanned crude extracts of *A. flavipes* CNL-338 for the presence of **9-11**, t**he** compounds were not detected. Based on the**se** data, we speculated that low production of **8** by *A. flavipes* CNL-338 could be impeding o**ur** ability to detect its conversion to the talaronoids.

To determine if **8** is indeed an intermediate **in** talaronoid biosynthesis, we biosynthetica**lly** prepared ^13^C-enriched **8** in *S. cerevisiae* using [1-^13^C]acetate. Labeled material was administered to *A. flavipes* CNL-338 and HREIMS inspection of the crude extract showed production of a new compound not observed in the DMSO control. The isotopic fragmentation pattern of the new compound also indicated it was derived from labeled material (Figure 4a). Closer inspection of the new compound showed that its retention time and *m/z* matched to that of an authentic standard of talarodiene C **11** (Figure 4b), thereby confirming that **8** had been biotransformed to **11**. Thus, the stable tracer isotope experiment confirmed **8** as an intermediate in the talaronoid biosynthetic pathway and established that TndB, TndE and TndF modify the 5-8-6 tricyclic hydrocarbon skeleton to afford talaronoid C.

**Figure 4.**
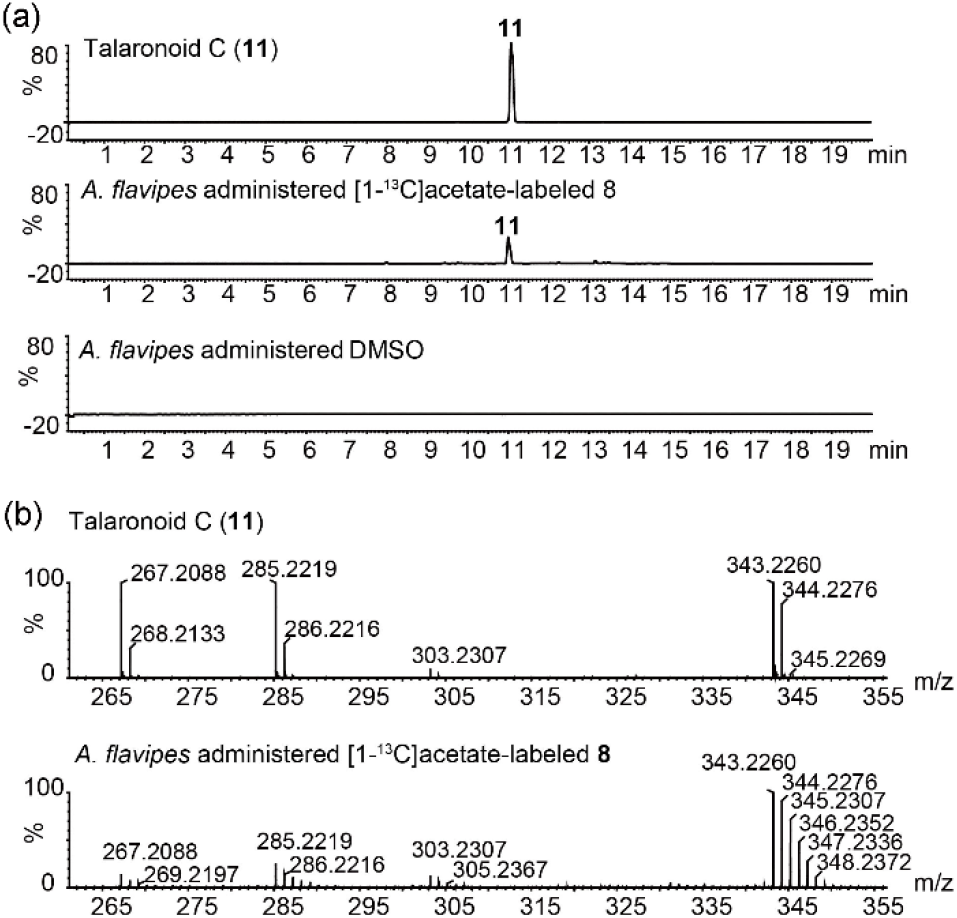
*In vivo* conversion of the talarodiene backbone **8** in *A. flavipes* CNL-338. (a) High-resolution LC-MS analysis (EIC = 343 *m/z*) for the [M+Na]^+^ adduct of the talaronoid C standard **11** compared to *A. flavipes* CNL-338 administered [1-^13^C]acetate-labeled **8** or a DMSO control. All traces are shown in the same scale. (b) HREIMS fragmentation pattern of the talaronoid C standard compared to the produ**ct** observed after administering [1-^13^C]acetate-labeled **8** to *A. flavipes* CNL-338.

In summary, we identified and characterized the *tnd* biosynthetic cluster responsible for the production of talaronoid C from the marine-derived fungus *A. flavipes* CNL-338. Heterologous expression of a cryptic type I bifunctional terpene synthase led to the discovery of a new diterpene possessing a benzo[*a*]cyclopenta[*d*]cyclooctane ring system and demonstrated that a single enzyme is responsible for the synthesis of this complex hydrocarbon scaffold. ^13^C-labeling studies helped elucidate a possible cyclization mechanism that would convert geranylgeranyl diphosphate to the 5-8-6 tricyclic hydrocarbon skeleton, and stable tracer isotope experiments validated **8** as an intermediate in talaronoid biosynthesis. Our work therefore brought to light the product of a cryptic terpene biosynthetic cluster, and information gleaned from the characterization of TndC can assist with future genome mining predictions.

## Acknowledgements

We thank Drs William Fenical (Scripps Institution of Oceanography) for *A. flavipes* CNL-338, Joseph Chappell (University of Kentucky) for *S. cerevisiae* ZXM144, Jack Skalicky (University of Utah) for helpful NMR advice, and John Alan Maschek (University of Utah) for GCMS assistance. S.C.H thanks the ALSAM foundation and ARUP laboratory for graduate research fellowships and J.M.W. thanks the Gordon and Betty Moore Foundation (GBMF7621) and the US National Institutes of Health (1R01AI155694) for support.

---

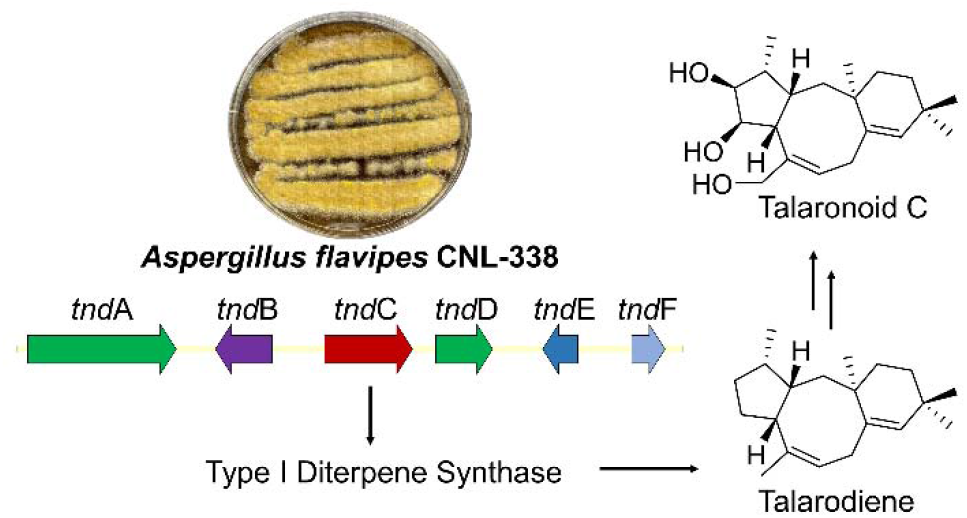

Our work identifying and characterizing a fungal bifunctional type I diterpene synthase expands the chemical space of terpene natural products, and our heterologous expression platform coupled with stable isotope tracer experiments presents a powerful platform for characterizing cryptic terpene biosynthetic clusters.

