## Supplemental for "Biochemical Characterization of a Fungal Bifunctional Type I Diterpene Synthase Brings a Cryptic Natural Product to Light"

### Table of Contents


**Pages**

**Experimental Procedures**

Strains S3
Chemicals and spectroscopic analysis S3
Culture conditions S3
General molecular biology procedures S3
Cloning and expression of TndC in *S. cerevisiae* S4
Constructing the *tndB* disruption cassette S4
Transformation of *tndB* disruption cassette into *A. flavipes* CNL-338 S4
Compound isolation and diterpene structure elucidation S5
Labeling studies with $^{13}C$-enriched precursors S5
Secondary metabolite analysis of wild-type *A. flavipes* CNL-338 and *ΔtndB* S5
*A. flavipes* CNL-338 biotransformation of $^{13}C$-enriched **8** S6
Accession number S6

**Supplementary Tables**

Table S1. Primers used in this study S7
Table S2. Spectral assignments of talarodiene **8** S8
Table S3. Percent enrichment of [1-$^{13}C$]acetate and [2-$^{13}C$]acetate- labeled **8** and coupling constants for [1,2-$^{13}C_2$]acetate-enriched **8** S9
Table S4. Annotation of the *tnd* gene cluster in *A. flavipes* CNL-338 S12

**Supplementary Figures**

Figure S1. Phylogenetic analysis of known fungal type I di/sesterterpenes S10
Figure S2. Terpene cyclase (TC) domain alignments S11
Figure S3. Alignment of the *tnd* gene cluster from *A. flavipes* CNL-338 and *T. stipitatus* S12
Figure S4. Alignment of tndC from the *tnd* cluster in *A. flavipes* and *T. stipitatus* S13
Figure S5. Alignment of tndB from the *tnd* cluster in *A. flavipes* and *T. stipitatus* S13
Figure S6. Reconstituting intron-free *tnd*C S14
Figure S7. High resolution LC-MS and GC-MS analysis of [1-$^{13}C$] acetate labeled **8** S15
Figure S8. Experimental and calculated ECD spectra of **8** S16
Figure S9. Disruption cassettes for the inactivation of *tnd*B and PCR verification S17
Figure S10. GC-MS comparison of **8** produced from *A. flavipes* and *⊗tndB* mutant S18
Figure S11. $^1H$ NMR spectrum of compound **8** in $CDCl_3$ S19
Figure S12. $^{13}C$ NMR spectrum of compound **8** in $CDCl_3$ S20
Figure S13. $^1H$-$^1H$ COSY spectrum of compound **8** in $CDCl_3$ S21
Figure S14. HSQCAD spectrum of compound **8** in $CDCl_3$ S22
Figure S15. HMBCAD spectrum of compound **8** in $CDCl_3$ S23
Figure S16. NOESY spectrum of compound **8** in $CDCl_3$ S24
Figure S17. ROESY spectrum of compound **8** in $CDCl_3$ S25
Figures S18–S22. 1D NOESY spectra of compound **8** in $CDCl_3$ S26–S30
Figure S23. $^{13}C$ NMR spectrum of [1-$^{13}C$]acetate-enriched **8** S31
Figure S24. $^{13}C$ NMR spectrum of [2-$^{13}C$]acetate-enriched **8** S32
Figure S25. $^{13}C$ NMR spectrum of [1,2-$^{13}C$]acetate-enriched **8** S33–S35

**Supplementary References** S36

### Experimental Procedures

#### *Strains*

The marine-derived fungal strain *Aspergillus flavipes* CNL-338 used in this study was kindly provided by Professor William Fenical at Scripps Institution of Oceanography. The strain was isolated as an endophyte from a red alga *Laurencia* sp. collected in the Bahamas,[1] and we recently reported the sequencing and assembly of its genome.[2] *Escherichia coli* DH5α and Top10 were used for routine cloning experiments. *Saccharomyces cerevisiae* ZXM144[3] (*sue, ∆erg9, ∆erg20, Mfps*) and the pXURA vector were kindly provided by Professor Joseph Chappell from the University of Kentucky and were used in this study for the *in vivo* characterization of the fungal type I diterpene synthase TndC.

#### *Chemicals and spectroscopic analysis*

Labeled [1-$^{13}$C]acetate and [2-$^{13}$C]acetate (99%) were purchased from Cambridge Isotope Laboratories, Inc. All solvents were HPLC grade from Fisher Scientific (Waltham, MA, USA). NMR spectra were obtained on a Varian INOVA 500 spectrometer with a 3 mm Nalorac MDBG probe. NMR conditions for the [1,2-$^{13}C_2$]acetate enrichment studies were as follows: spectral width, 31421.8 Hz ; pulse width, 9.5000 µsec; acquisition time, 0.2607 sec; and 224 accumulations. LC-MS analysis was performed on an Acquity Arc UHPLC/MS (Waters, Milford, MA, USA) in positive mode with a Waters XBridge C18 column (2.1 mm x 50 mm, 3.5 ⌠m). High resolution LC-MS was performed on an Acquity H-Waters Xevo G2-XS-Q-ToF (Waters) using an Acquity UPLC BEH C18 column (2.1 mm x 150 mm, 1.7 µm) with a linear gradient of 5–100% solvent B over 17 min (solvent A, $H_2O$ with 0.05% formic acid; solvent B, MeCN), and 1 µl aliquot of each sample was injected for analysis. Gas chromatography was performed on an Agilent HP 6890 GC Systems connected to a 5973 mass selective detector and an Agilent DB-5MS column (30 m x 0.25 mm x 0.25 µm). The temperature of the ionization chamber was 260 °C with electron impact ionization at 70 eV. The temperature of the column was held at 100 °C for three min, increased at a rate of 15 °C/min to 330 °C, and then held at 330 °C for 15 min. A 1 µl aliquot of each sample was injected for analysis. ECD spectra were taken on a JASCO J-815 CD spectrometer.

#### *Culture conditions*

For genomic DNA extraction, wild-type *A. flavipes* CNL-338 and the Δ*tnd*B mutant were grown in 3 mL YPM media (0.2% yeast extract, 0.2% peptone, 0.4% mannitol) containing 3.3% artificial sea salt (Instant Ocean, USA) in 10 x 60 mm petri dishes for three days at 30 °C under static conditions. For culturing the Δ*tnd*B mutant, 300 ⌠g/mL of zeocin was added for antibiotic selection. For RNA extraction, *A. flavipes* CNL-338 was cultivated in 30 mL YPG liquid media (1% dextrose, 0.5% peptone, and 0.5% yeast extract) containing 3.3% artificial sea salt in 100 mL Erlenmeyer flasks for three days at 30 °C and 180 rpm. Transformants of *S. cerevisiae* ZXM144 were cultured on synthetic complete dropout media supplemented with ergosterol (SCE) (0.6% succinic acid, 0.5% potassium hydroxide, 2% dextrose, pH 5.3). After sterilization, 10% nitrogen base solution (1.7 % yeast nitrogen base without amino acids and 5% ammonium sulfate) was added by filter sterilization, as well as 0.14% amino acid dropout powder (Sigma yeast synthetic drop-out media supplement without Leu, Trp, Ura, and His), 0.025% His, 0.02% Met, 0.015% Trp, either 0.1% Leu or 0.03% Ura, and 0.4% (v/v) ergosterol (10 mg/mL ergosterol dissolved in 1:1 Triton X-100:ethanol). Positive colonies were cultured in large-scale YPDE liquid media (1% yeast extract, 2% peptone, 2% dextrose and 0.4% (v/v) ergosterol stock solution pH 5.3).

#### *General molecular biology procedures*

PCR reactions were carried out using AccuPrime Taq DNA polymerase (Invitrogen). PCR screening of transformants was performed using GoTaq® Master Mix (Promega). All restriction enzymes were purchased from New England Biolabs. Primers were synthesized by Integrated

DNA Technologies (Coralville, IA, USA). pCR™-blunt (Invitrogen) was used to construct recombinant DNA products, and DNA sequencing was performed by GeneWiz (South Plainfield, NJ, USA). Oligonucleotide sequences used in this study are provided in Table S1. RNA was extracted from a three-day old YPG culture using the RiboPure Yeast kit (Ambion) following the manufacturer's instructions, and contaminating gDNA was digested with DNase I (2 U/µL) (Invitrogen) at 37 °C for four hours. Complementary DNA (cDNA) was synthesized from total RNA using the Reverse Transcription Kit (Promega) according to the manufacturer's protocol using the Anchored Oligo$(dT)_{20}$ primer (Invitrogen).

***Cloning and expression of TndC in S. cerevisiae strains***

For heterologous expression of the bifunctional diterpene synthase TndC in *S. cerevisiae* ZXM144, intron-free sequences of *tnd*C were amplified from cDNA. Briefly, primer pairs 338_diTS_P1_F with 338_diTS_P1_R and 338_diTS_P2_F with 338_diTS_P2_R were used to amplify two overlapping pieces for the reconstitution of *tnd*C. The amplified fragments were subcloned into pCR®-blunt (Invitrogen) according to the manufacturer's protocol and sequence verified. Intron-free *tnd*C was then amplified using primer pairs 338_diTS_pXURA_mFPS144_F and 338_diTS_pXURA_mFPS144_R, digested with XmaI and XhoI, and cloned into the XmaI-XhoI sites of pXURA-*pGPD::mFPS144* under the *TEF* promoter to create the plasmid pPZW01 (Figure S6). The plasmid was then transformed into *S. cerevisiae* ZXM144 competent cells using the LiOAc-PEG method. Briefly, a single yeast colony was used to inoculate 3 mL YPDE and cultured at 30 °C with 200 rpm shaking for about 30 hours. A 0.5 mL aliquot from each 3 mL culture was then used to inoculate 50 mL YPDE liquid media in 500 mL Erlenmeyer flasks and cultured at 30°C with constant shaking at 200 rpm for 12–15 hours. The cells were then pelleted by centrifugation in sterile 50 mL Falcon tubes at 3738 x g for ten minutes at 4 °C. The pelleted cells were re-suspended in 25 mL of 1.2 M sorbitol wash buffer (1.2 M sorbitol in 0.1 M potassium phosphate buffer, pH 7.5) by gently inverting and swirling the tube. Competent cells were prepared by re-pelleting and re-suspending the cells in 3.55 mL of LiOAc-PEG transformation buffer (0.1 M LiOAc, 33% PEG $M_n$ 3350 and 280 µg/mL single stranded salmon sperm DNA). For each transformation, 1–5 µg of plasmid DNA was added to 200 µL of competent cells, incubated at 30 °C for one hour, and vortexed for one sec every 15 min. The cells were then heat shocked at 42 °C for ten minutes. Afterward, they were plated on SCE dropout medium and incubated at 30 °C for 3–5 days.

***Constructing the tndB disruption cassette***

To confirm the function of TndB in talaronoid biosynthesis, the *tnd*B gene was targeted for gene inactivation using a disruption cassette (Figure S9). The inactivation cassette was designed to contain the zeocin resistance gene *Sh ble* and the constitutive promoter PtrpC.[2] Fragments of approximately 1 kb both upstream and downstream of *tnd*B were amplified from *A. flavipes* CNL-338 genomic DNA using primer pairs P450_5F_F with P450_5F_R and P450_3F_F with P450_3F_R, respectively. The zeocin resistance marker was amplified using primer pairs PtrpC fwd with zeocin R. The three PCR fragments were fused together using fusion PCR[4] to generate the *tnd*B disruption cassette (Figure S9A). The disruption cassette was ligated into pCR®-blunt for DNA sequencing before being amplified using primer pairs P450_5F_F with P450_3F_R.

***Transformation of tndB disruption cassette into A. flavipes CNL-338***

The PCR amplified linear *tnd*B disruption cassette (~ 10 µg) was transformed into *A. flavipes* CNL-338 protoplasts, as described previously[5]. Transformants were grown on stabilized minimal agar medium (1.2 M sorbitol, 1.5% agar, 1% dextrose, 5% nitrate salts, 0.1% trace elements) supplemented with 300 µg/mL zeocin. For verification of correct integration into the genome, gDNA was extracted and used as a template for PCR using primer pairs P450_pESCleu2_F with P450_pESC-leu2_R, P450_5F_Check_F with PtrpC rev screen, and zeo fwd screen with P450_3F_Check_R (Figure S9).

***Compound isolation and diterpene structure elucidation***

For the isolation of the diterpene **8**, *S. cerevisiae* ZXM144 harboring the plasmid pPZW01 was cultured in 3 mL $SCE_{glu}$(-Ura) liquid media for five days at 23 °C with constant shaking at 250 rpm. The 3 mL seed culture was then used to inoculate 5 x 1 L cultures of YPDE in 2.8 L Fernbach flasks and allowed to shake at 23°C for seven days at 250 rpm. The 5 L culture was extracted with 5 L acetone and partitioned with 10 L hexanes. The hexanes phase was concentrated *in vacuo* to yield a 2.4 g crude extract. The crude extract was then re-dissolved in hexanes and fractionated using a normal-phase Silica gel column and eluting with 100% hexanes to afford fractions 1–30. The different fractions (F1–F30) were first analyzed by thin layer chromatography (TLC) developed with 100% hexanes and stained with vanillin (10 mL ethanol with 0.6 g vanillin and 200 µL $H_2SO_4$,). TLC showed the target compound **8** was located in fraction seven. To further purify compound **8**, fraction F7 was subjected to preparative TLC developed with 100% hexanes to afford pure compound **8** (6 mg) as a colorless oil. Compound **8** was dissolved in $CDCl_3$ and its chemical structure was elucidated by 1D and 2D NMR (Figures S11–S22 and Table S2). High resolution LC-MS indicated the molecular weight was 273.2200 $[M+H]^+$ (Figure S7A), and GC-MS confirmed the fragmentation pattern was characteristic of a terpene (Figure S7B). The absolute configuration of **8** was determined by ECD experiments and calculations. The time-dependent density-functional theory (TD-DFT)-calculated ECD spectra of **8i** (2*S*,3*S*,6*R*,11*R*) and **8ii** (2*R*,3*R*,6*S*,11*S*) were obtained at the B3LYP/6-31+(d,p) level with MeOH as solvent. The data showed that the experimental ECD spectrum of **8** was consistent with the calculated ECD of **8i** (Figure S8).

***Labeling studies with $^{13}$C-enriched precursors***

For the stable isotope enrichment studies, 200 mg of [1-$^{13}$C]acetate, [2-$^{13}$C]acetate or [1,2-$^{13}C_2$]acetate was dissolved in 1 mL Milli-Q water and filter sterilized. At the time of inoculation, a 1 mL aliquot of each isotope was independently added to a 1 L YPDE culture of *S. cerevisiae* ZXM144 transformed with the pPZW01 plasmid in 2.8 L Fernbach flasks. The cultures were grown at 23 °C with constant shaking at 250 rpm for seven days. The 1 L cultures were extracted with 1 L acetone and partitioned with 2 L hexanes. The isolation of labeled **8** from the respective [1-$^{13}$C]acetate-enriched crude extract (560 mg), [2-$^{13}$C]acetate-enriched crude extract (570 mg) or [1,2-$^{13}C_2$]acetate-enriched crude extract (554 mg) followed the same procedures as described above. $^{13}$C NMR data of [1-$^{13}$C]acetate-, [2-$^{13}$C]acetate- and [1,2-$^{13}C_2$]acetate-enriched **8** were collected in $CD_3Cl$. The percent incorporation for the [1-$^{13}$C] and [2-$^{13}$C], as well as one bond homonuclear $^{13}$C,$^{13}$C scalar couplings, were calculated using established methods.[6]

***Secondary metabolite analysis of wild-type* A. flavipes *CNL-338 and Δ*tnd*B***

Wild-type *A. flavipes* CNL-338 and the mutant Δ*tnd*B strain were cultured in 30 mL of YPG liquid medium supplemented with 3.3% artificial sea salt in 250 mL Erlenmeyer flasks with constant shaking at 180 rpm at 30 °C for 9 days. The cultures were extracted three times with equal volume of ethyl acetate. The ethyl acetate fractions were combined and concentrated *in vacuo* to yield ~ 250 mg crude extracts. Each crude extract was dissolved in 1 mL HPLC grade ethyl acetate, and a 1 µl aliquot of each sample was injected for analysis by GC-MS using the parameters described above (Figure S10).

#### *A. flavipes* CNL-338 biotransformation of $^{13}C$-enriched 8

Wild-type *A. flavipes* CNL-338 was cultivated in 2 x 250 mL Erlenmeyer flasks containing 30 mL YPG liquid medium with 3.3% artificial sea salt at 30 °C with constant shaking at 180 rpm for seven days. At the time of inoculation and every day for five days, [1-$^{13}C$]acetate-enriched **8** (400 µg in 40 µL DMSO) was added to one of the 30 mL cultures. A 40 µL aliquot of DMSO was continuously added to the other culture for five days as a control. In total, over the five-day fermentation, 2 mg of labeled **8** was added. On the seventh day, each culture was extracted three times with equal volumes of ethyl acetate, and the ethyl acetate extracts were combined and concentrated *in vacuo* to yield the ethyl acetate crude extract. The [1-$^{13}C$]acetate-enriched **8** provided a 313 mg crude extract, while the crude extract from the DMSO control experiment was 320 mg. Each extract was resuspended in 500 ⌠L methanol. The talaronoid C **11** standard, and the extracts of wild-type *A. flavipes* CNL-338 administered DMSO, or **8** labeled with [1-$^{13}C$]acetate were analyzed by high resolution LC-MS on an Acquity H-Waters Xevo G2-XS-Q-ToF (Waters) using an Acquity UPLC BEH C18 column (2.1 mm x 150 mm, 1.7 µm, 0.3 mL/min) with a linear gradient of 5–100% solvent B over 17 min (solvent A, $H_2O$ with 0.05% formic acid; solvent B, MeCN), and a 1 µl aliquot of each sample was injected for analysis (Figure 5).

#### Accession number

The sequence of the *tnd* biosynthetic gene cluster has been deposited in the GenBank database (NCBI) with the accession number MW248390.

### Supplementary Results

***Supplementary Table S1.*** Primers used in this study

| Primer | Sequence shown 5’–3’ |
|---|---|
| 338_diTS_P1_F | TCAACTATCAACTATTAACTATATCGTAATACCATATGGAATACAGGTACTC TACGG |
| 338_diTS_P1_R | GATGAACTCCATTGCCATTGG |
| 338_diTS_P2_F | GCTTGTTGTACAGCCTGAGC |
| 338_diTS_P2_R | TGTCATTTAAATTAGTGATGGTGATGGTGATGCACGGCCTTCCGCAGCAT CTC |
| 338_diTS_pXURA_mFPS144_F | GG**CCCGGG**ATGGAATACAGGTACTCTACGG |
| 338_diTS_pXURA_mFPS144_R | TA**CTCGAG**TTAGTGATGGTGATGGTGATGC |
| P450_5F_Check_F | CCTCGGCTGGATTGTGATG |
| P450_5F_F | CAGCCACATAACTTGCAACG |
| P450_5F_R | CTCCTTCAATATCATCTTCTGTCGACCGGCTGTAAGGGACGACTT |
| PtrpC fwd | GTCGACAGAAGATGATATTGA |
| zeocin R | AGCTTGCAAATTAAAGCCTT |
| PtrpC rev screen | TGTCCTCGTTCCTGTCTGC |
| zeo fwd screen | TTCGTGGAGGACGACTTCG |
| P450_3F_F | GCTCGAAGGCTTTAATTTGCAAGCTGTGGATGTGCTGGATGTGG |
| P450_3F_R | CAATGCCATGCACGCTTTG |
| P450_3F_Check_R | CGTCTGGTTGGTGGTGAAC |

Note: XmaI and XhoI restriction recognition sites are bolded and underlined.

***Supplementary Table S2:*** NMR assignments for talarodiene **8** ($^{1}$H NMR at 500 MHz; $^{13}$C NMR at 125 MHz in $CDCl_3$). Numbering of carbon atoms is based on the geranylgeranyl diphosphate precursor.

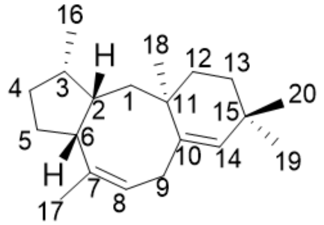


Talarodiene (**8**)

| Carbon | $\delta_C$ (ppm) | $\delta_H$ (*J* in Hz) | $^{1}$H-$^{1}$H COSY | HMBC | NOESY |
|---|---|---|---|---|---|
| 1 | 39.3 $CH_2$ | 1.80, t (13.3); 1.19, m (overlap) | 2 | | |
| 2 | 45.5 CH | 2.20, m | 1, 3, 6 | | 6, 20 |
| 3 | 36.6 CH | 2.05, m | 2, 4, 16 | | |
| 4 | 33.5 $CH_2$ | 1.26, m; 1.95, m | 3, 5 | | |
| 5 | 28.5 $CH_2$ | 1.87, m; 1.52, m | 4, 6 | | |
| 6 | 39.9 CH | 3.34, t (8.1) | 2, 5 | | 2, 20 |
| 7 | 134.9 C | | | | |
| 8 | 125.1 CH | 5.35, brs | 9 | | |
| 9 | 37.1 $CH_2$ | 2.94, dd (6.5, 19.4); 2.68, br d (18.2) | 8 | 7, 10, 14 | |
| 10 | 141.7 C | | | | |
| 11 | 38.0 C | | | | |
| 12 | 33.5 $CH_2$ | 1.20, m; 1.94, m | 13 | | |
| 13 | 34.2 $CH_2$ | 1.50, m; 1.42, m | 12 | | |
| 14 | 135.0 CH | 5.15, s | | | |
| 15 | 32.4 C | | | | |
| 16 | 18.2 $CH_3$ | 0.78, d(7.4) | 3 | 2, 3, 4 | 18 |
| 17 | 23.3 $CH_3$ | 1.74, s | | 6, 7, 8 | |
| 18 | 29.4 $CH_3$ | 1.08, s | | 1, 10, 11, 12 | 16 |
| 19 | 28.3 $CH_2$ | 0.97, s | | 14, 15 | |
| 20 | 31.3 $CH_3$ | 0.95, s | | 15, 13 | 2, 6 |

***Supplementary Table S3:*** $^{13}$C and $^{1}$H assignments, percent incorporation from [1-$^{13}$C]acetate and [2-$^{13}$C]acetate enrichment experiments, and coupling constants for [1,2-$^{13}C_2$]acetate-enriched **8**. Numbering of carbon atoms is based on the geranylgeranyl diphosphate precursor.

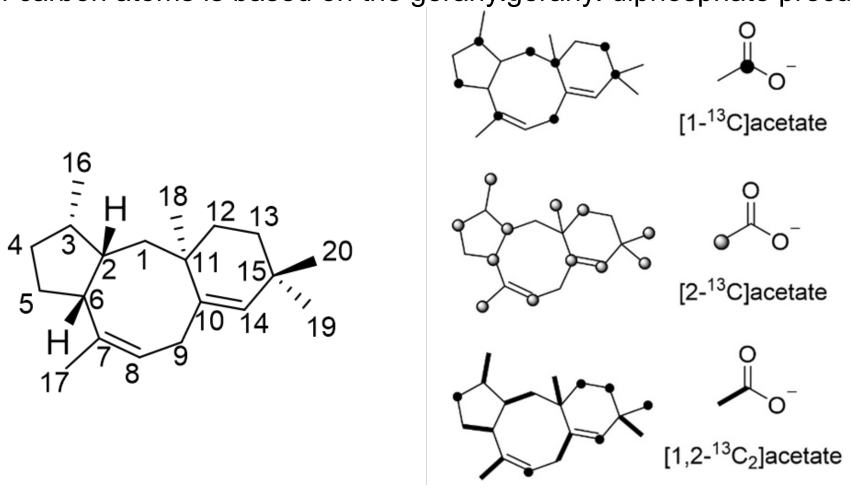


| Carbon | | Incorporation (%) | | [1,2-$^{13}C_2$]acetate |
|---|---|---|---|---|
| | $\delta_C$ [a] | [1-$^{13}$C]-sodium acetate[b,c] | [2-$^{13}$C]-sodium acetate[b,d] | $J_{CC}$(Hz) |
| 1 | 39.3 $CH_2$ | 7.3 | | 38 |
| 2 | 45.5 CH | | 21.2 | 38 |
| 3 | 36.6 CH | 10.6 | | 35 |
| 4 | 33.5 $CH_2$ | | 26.4 | S |
| 5 | 28.5 $CH_2$ | 11.7 | | 34 |
| 6 | 39.9 CH | | 19.8 | 34 |
| 7 | 134.9 C | 44.7 | | 44 |
| 8 | 125.1 CH | | 36.3 | S |
| 9 | 37.1 $CH_2$ | 9.4 | | 43 |
| 10 | 141.7 C | | 33 | 43 |
| 11 | 38.0 C | 8.6 | | 35 |
| 12 | 33.5 $CH_2$ | | 29.2 | S |
| 13[d] | 34.2 $CH_2$ | 8.5 | | S |
| 14 | 135.0 CH | | 36 | S |
| 15 | 32.4 C | 0.75 | | 35 |
| 16 | 18.2 $CH_3$ | | 29 | 35 |
| 17[c] | 23.3 $CH_3$ | | 25.3 | 44 |
| 18 | 29.4 $CH_3$ | | 22.3 | 35 |
| 19 | 28.3 $CH_2$ | | 22.7 | 35 |
| 20 | 31.3 $CH_3$ | | 23.4 | S |

[a] $^{13}$C NMR spectral data taken at 125 MHz in $CDCl_3$; [b] Percent incorporation (A - B)/B; A = intensity of enriched carbon and B = intensity of natural abundance carbon; [c] Incorporation relative to the natural abundance carbon at 23.3 ppm (C17); [d] Incorporation relative to natural abundance carbon at 34.2 ppm (C13). S = singly enriched

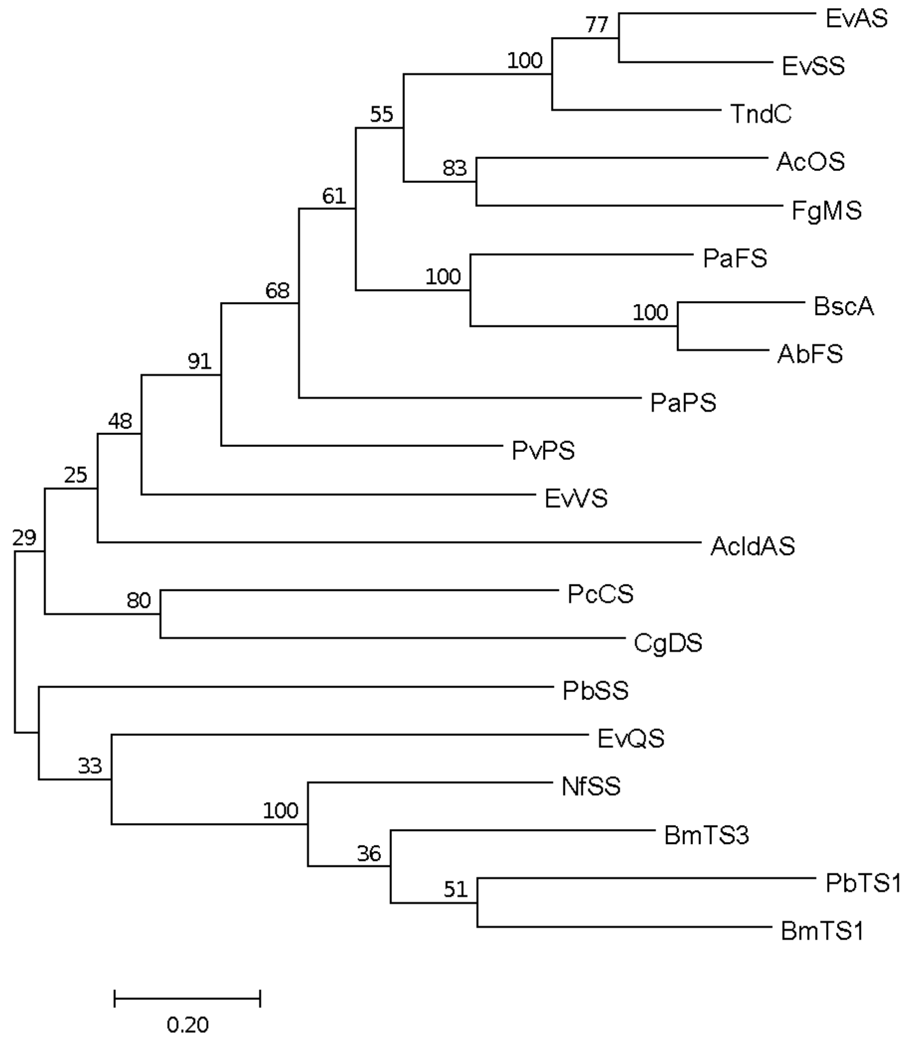


***Supplementary Figure S1.*** Relatedness of fungal type I di/sesterterpenes with TndC. Multi-sequence alignment using characterized bifunctional fungal terpene synthases and TndC was performed using MUSCLE,[7] and the unrooted Maximum Likelihood tree was created in Mega.[8] The scale bar indicates 0.20 changes per amino acid. Accession numbers of sequences used to create the phylogenetic tree are PaFS (XM_036618969.1),[9] AcOS (ACLA_076850),[10] NfSS (NFIA_055500),[11] PbTS1 (LC274619 in DNA Data Bank of Japan (DDBJ)),[12] FgMS (FgJ07578),[13] EvAS (LC113889),[14] EvQS (LC155210),[15] EvSS (LC073704.1),[16] EvVS (LC063849.1),[17] PvPS (LC228602.1), PaPS (UniProtKB/Swiss-Prot: P9WEV6.1), PcCS (LC411963.1),[18] PbSS (LC228601.1),[19] AbFS (AB465604), [20] AcldAS (CEL06489.1),[21] BmTS1 (XM_014217355.1),[12] BmTS3 (XM_014217731.1),[12] BscA (XP_007924161.1),[22] and CgDS (CGL13742.1).[23]

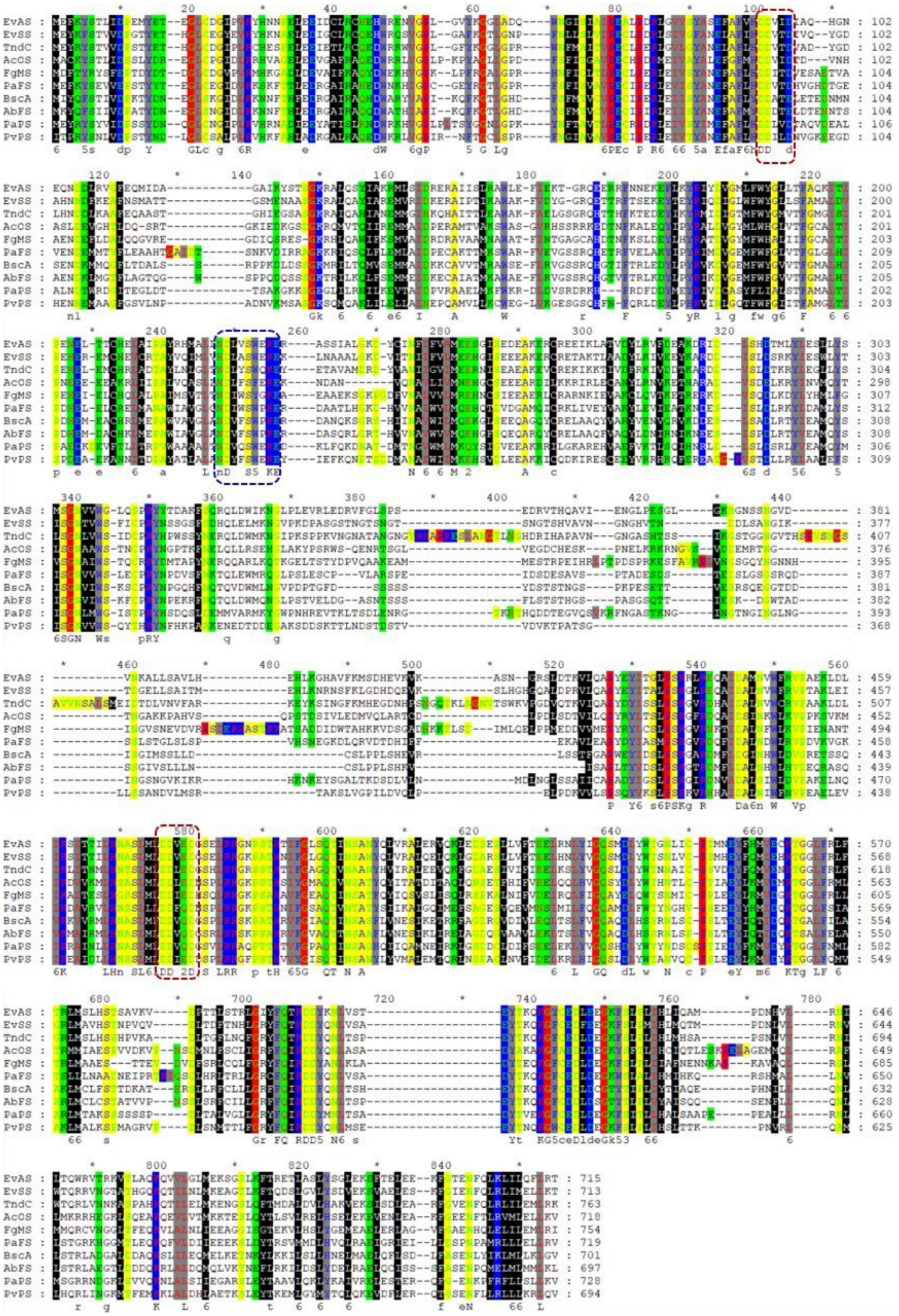

***Supplementary Figure S2.*** Alignment of TndC and selected fungal bifunctional terpene synthases. The conserved DDXXD/E (in TC domain) and DDXXD/N (in PT domain) motifs and the NSE/DTE motif are marked by red and blue dashed boxes, respectively.

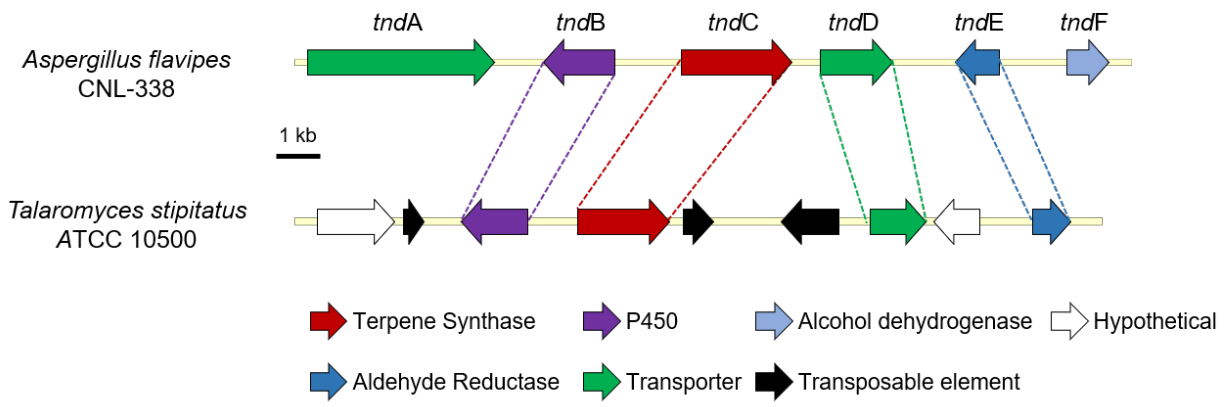


***Supplementary Figure S3.*** Alignment of the *tnd* gene cluster from *Aspergillus flavipes* CNL-338 and *Talaromyces stipitatus* ATCC 10500.

***Supplementary Table 4.*** Annotation of the *tnd* gene cluster in *Aspergillus flavipes* CNL-338

| **Gene product** | **Proposed function** | **Sequence similarity (origin)** | **Coverage/ identity (%)** | **Accession number** |
|---|---|---|---|---|
| *tnd*A | Pleiotropic drug resistance proteins (PDR1-15), ABC superfamily | *Penicillium brasilianum* | 99/86 | CEJ59027 |
| *tnd*B | Cytochrome P450 | *Talaromyces stipitatus ATCC 10500* | 100/80 | TSTA_007840 |
| *tnd*C | Geranylgeranyl diphosphate synthase/talarodiene synthase | *Talaromyces stipitatus ATCC 10500* | 99/78 | TSTA_007830A |
| *tnd*D | MFS multidrug transporter, putative | *Talaromyces stipitatus ATCC 10500* | 90/81 | TSTA_007800 |
| *tnd*E | Aldehyde reductase | *Talaromyces stipitatus ATCC 10500* | 87/67 | TSTA_007780 |
| *tnd*F | Alcohol dehydrogenase | *Aspergillus sclerotialis* | 99/85 | RJE24497 |

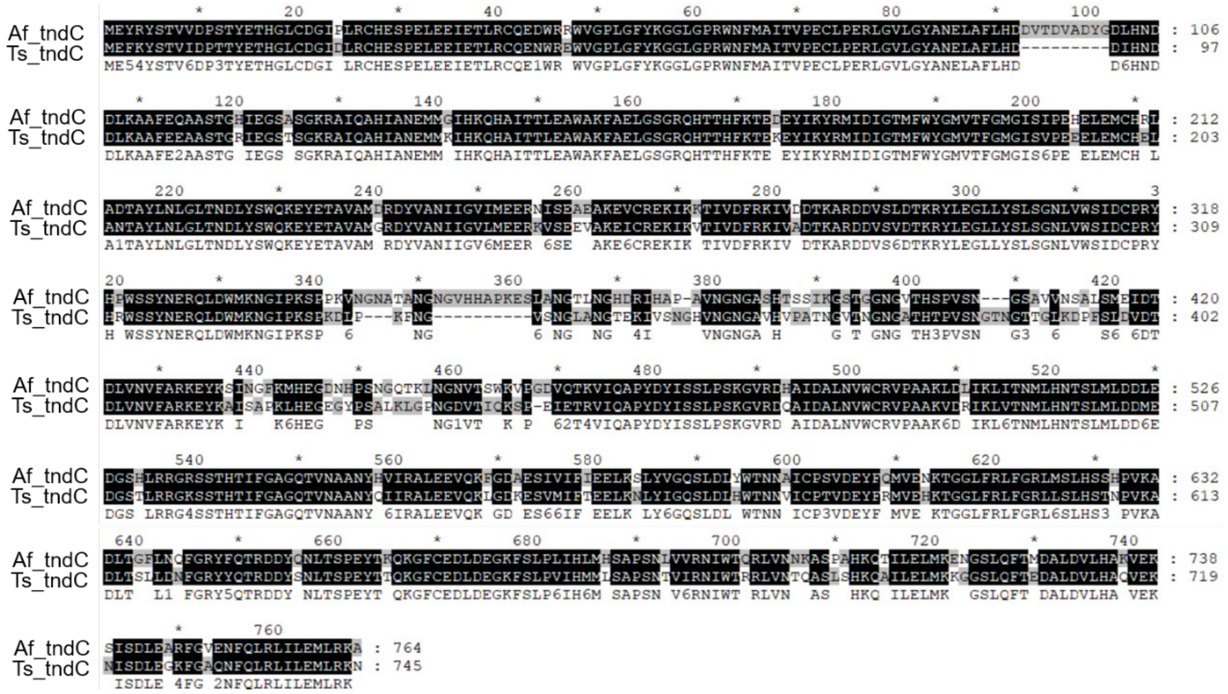


***Supplementary Figure S4.*** Alignment of the diterpene synthase *tnd*C from the *tnd* cluster in *Aspergillus flavipes* CNL-338 and *Talaromyces stipitatus* ATCC 10500.

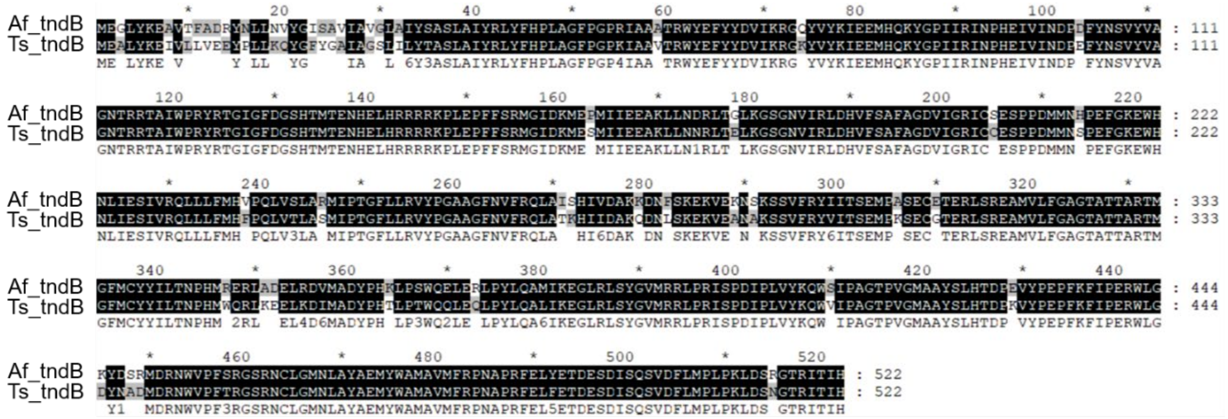


***Supplementary Figure S5.*** Alignment of the P450-*tnd*B from the *tnd* cluster in *Aspergillus flavipes* CNL-338 and *Talaromyces stipitatus* ATCC 10500.

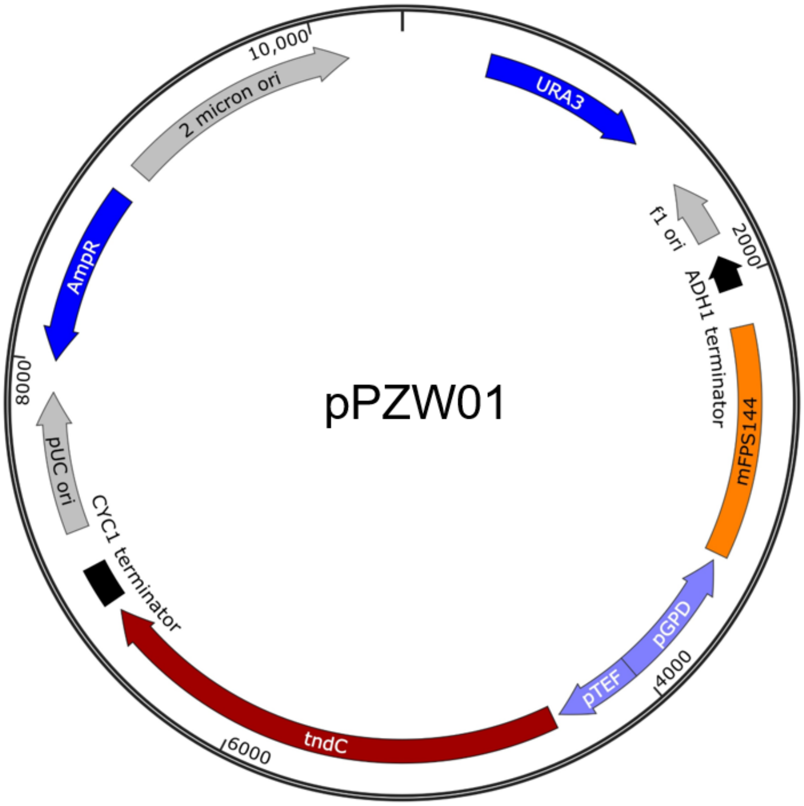


***Supplementary Figure S6.*** Reconstituted intron-free *tnd*C. The bifunctional diterpene synthase *tnd*C was inserted into the XmaI-XhoI sites of pXURA-GPD::mFPS144 under the constitutive pTEF promoter to yield the pPZW01 plasmid.

A)

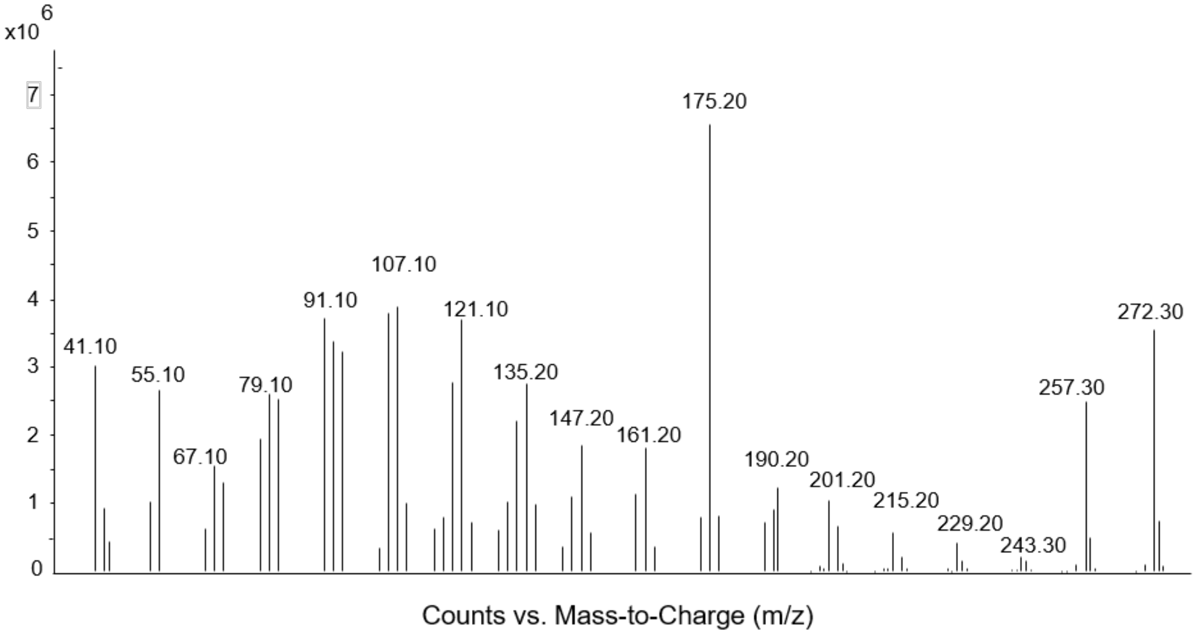


B)

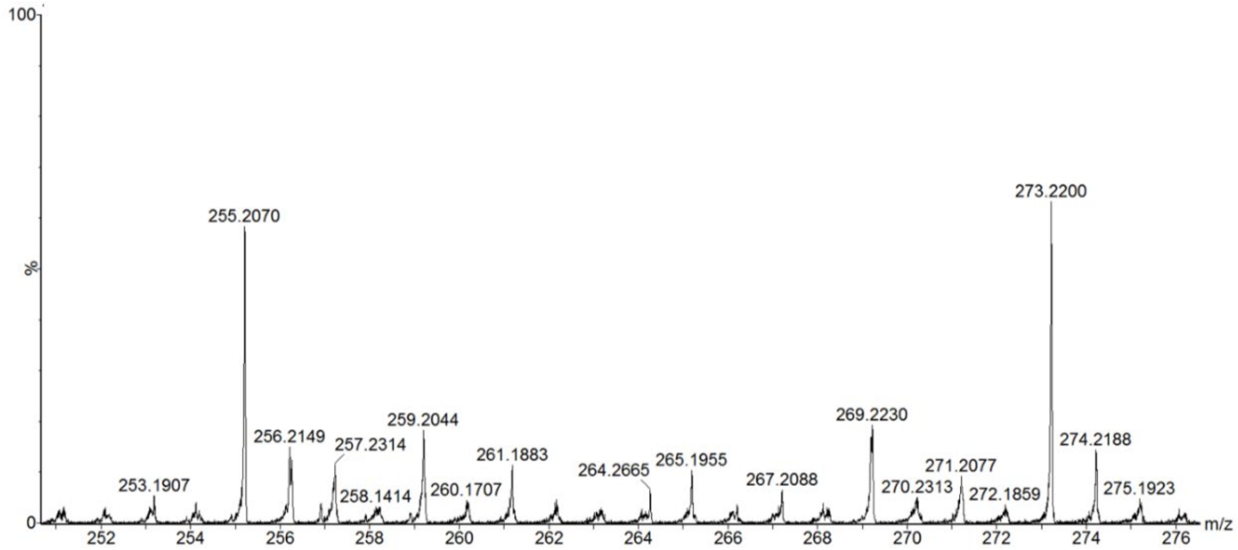


***Supplementary Figure S7.*** GC-MS fragmentation pattern of **8** (A) and high resolution LC-MS analysis of **8** (B).

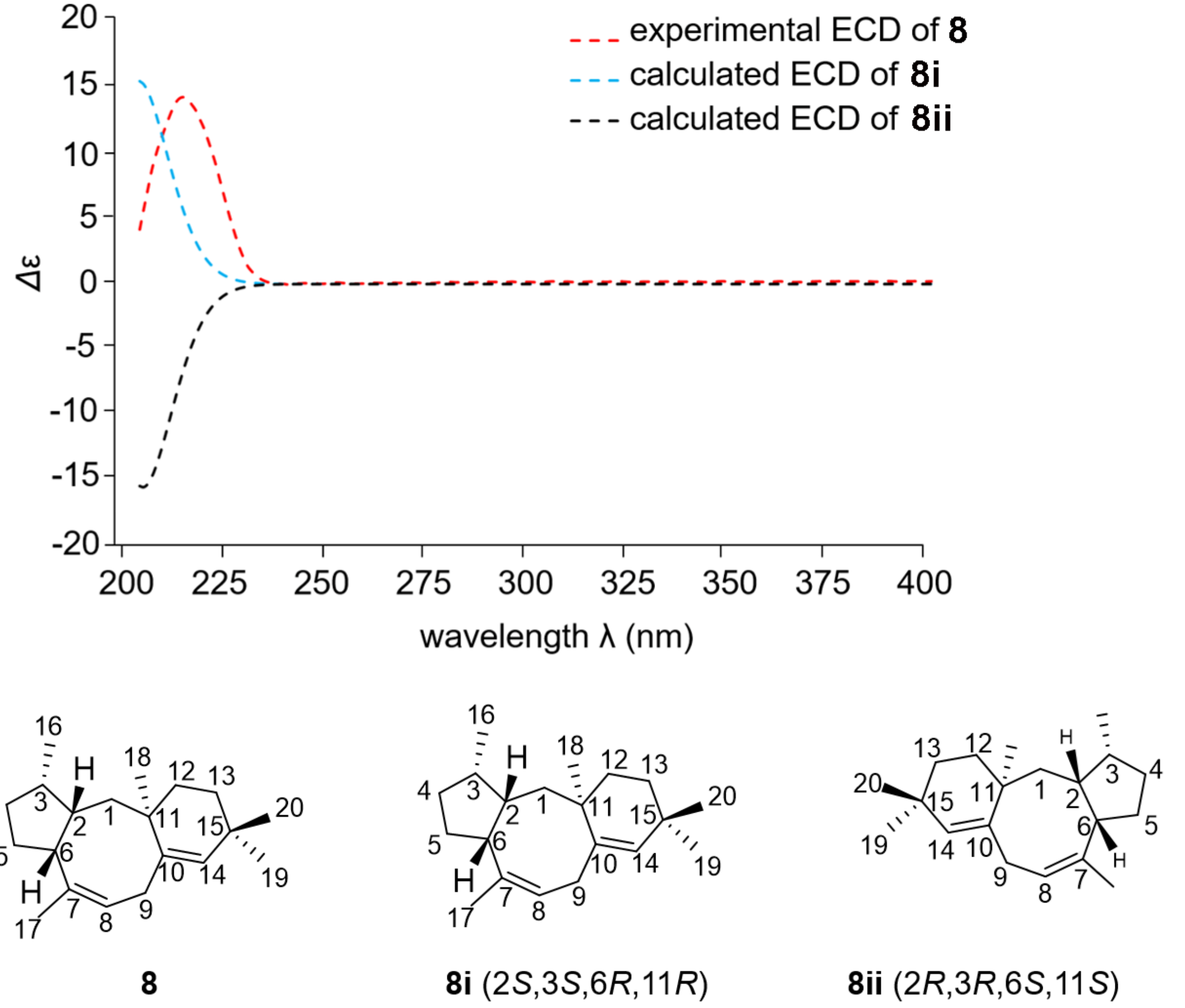


***Supplementary Figure S8.*** Experimental and calculated ECD spectra of **8**.

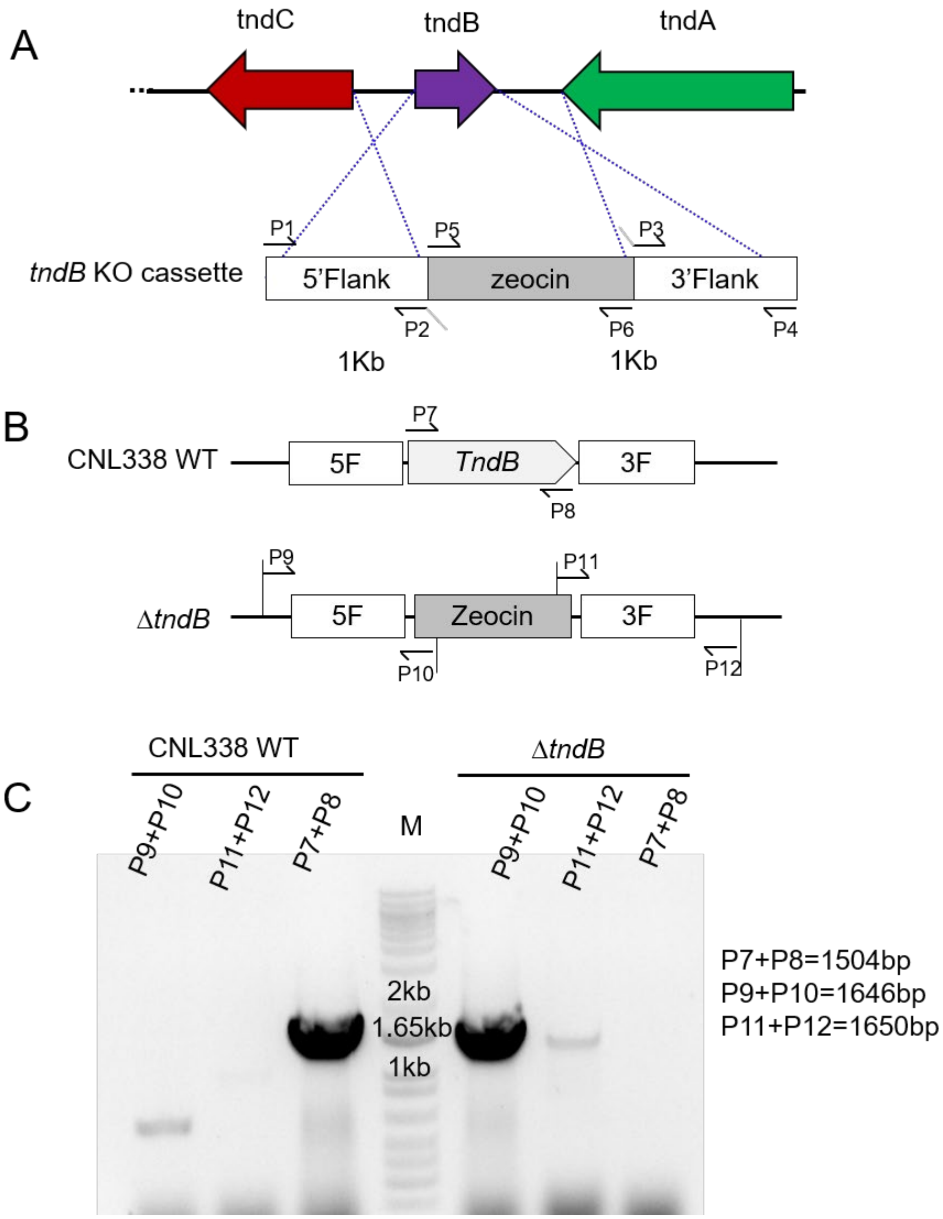


***Supplementary Figure S9.*** Creating the disruption cassette for the inactivation of *tnd*B in *A. flavipes* CNL-338 and PCR verification of its integration in the genome. Primers to screen for the presence and integration of the disruption cassette include P450_5F_F (P1), P450_5F_R (P2), P450_3F_F (P3), P450_3F_R (P4), PtrpC fwd (P5), zeocin R (P6), P450_pESC-leu2_F (P7), P450_pESC-leu2_R (P8), P450_5F_Check_F (P9), PtrpC rev screen (P10), zeo fwd screen (P11) and P450_3F_Check_R (P12).

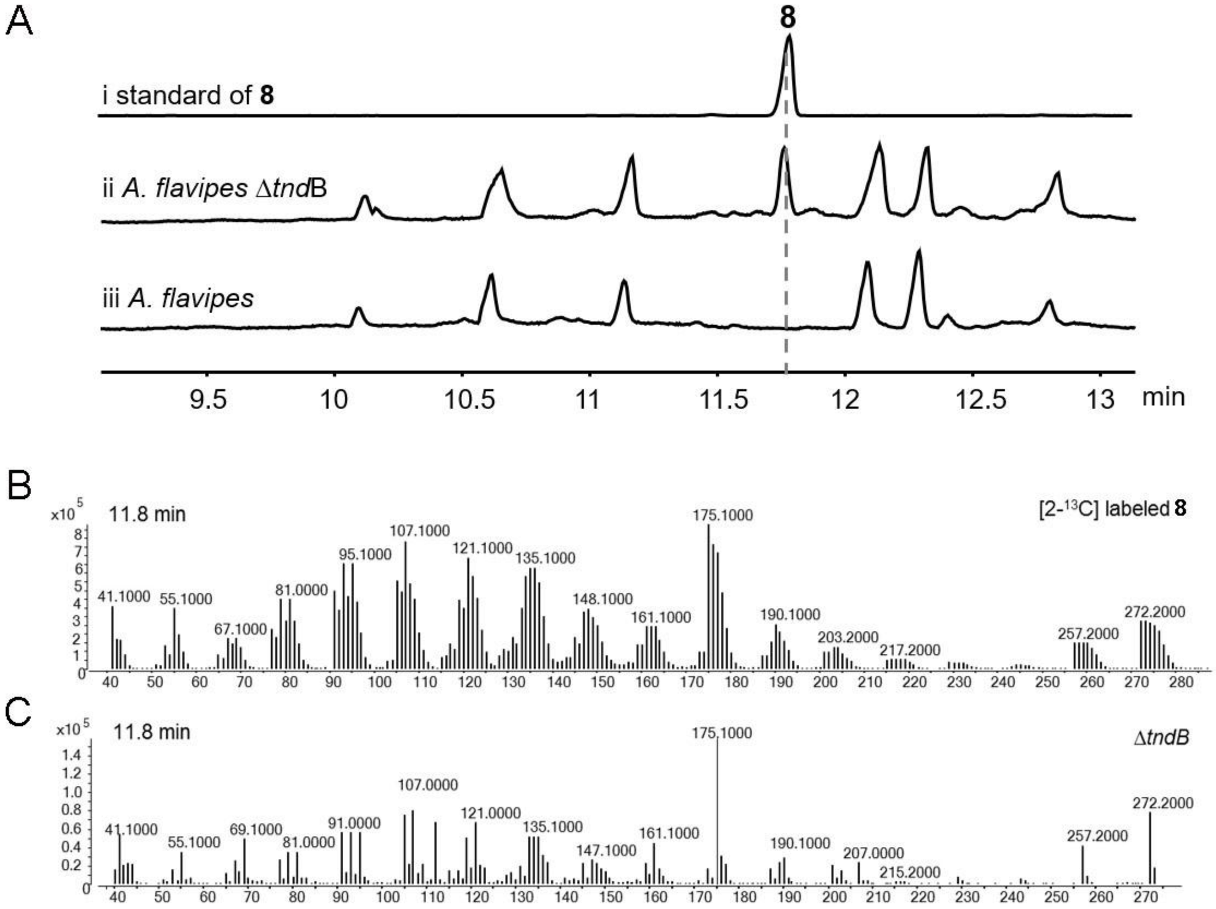


***Supplementary Figure S10.*** GC-MS comparison of **8** produced from *A. flavipes* CNL-338 and the Δ*tnd*B mutant. (A) GC-MS chromatogram of (i) [2-$^{13}$C]acetate-enriched **8** isolated from *S. cerevisiae* ZXM144 harboring a plasmid-borne *tnd*C*;* (ii) crude extract from *A. flavipes* CNL-338 wide-type; and (iii) Δ*tnd*B mutant strain. (B) GC-MS fragmentation pattern of [2-$^{13}$C]acetate-enriched **8** (trace i in panel A with a retention time of 11.8 min). (C) GC-MS of **8** identified in the Δ*tnd*B mutant (trace iii in panel A with a retention time of 11.8 min). The dominant mass fragments at m/z 272, 257, 190, 175, and 161 match with the mass fragments from [2-$^{13}$C]-labeled **8** shown in panel B.

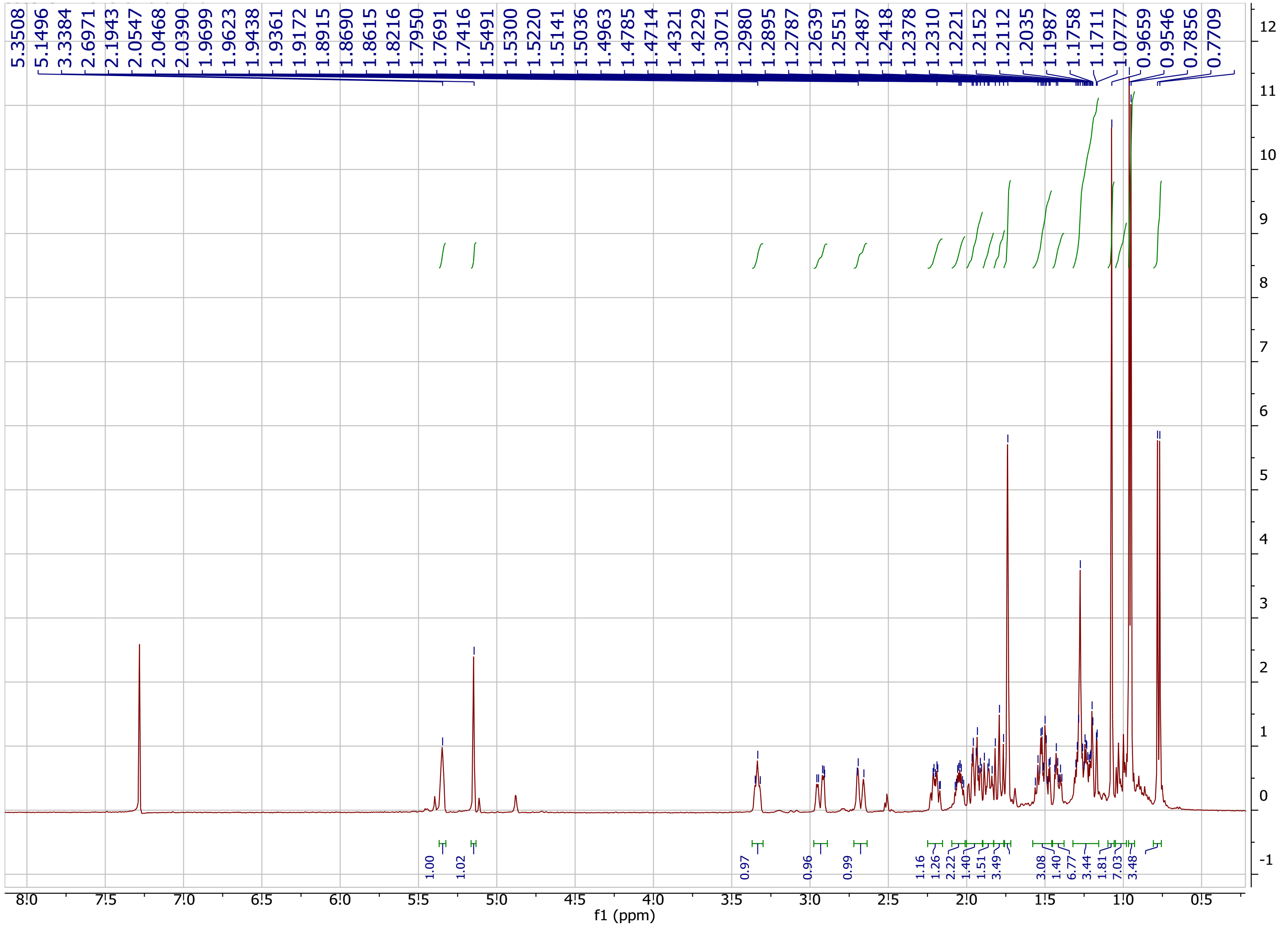

***Supplementary Figure S11***. $^{1}$H NMR spectrum (500 MHz) of compound **8** in $CDCl_3$.

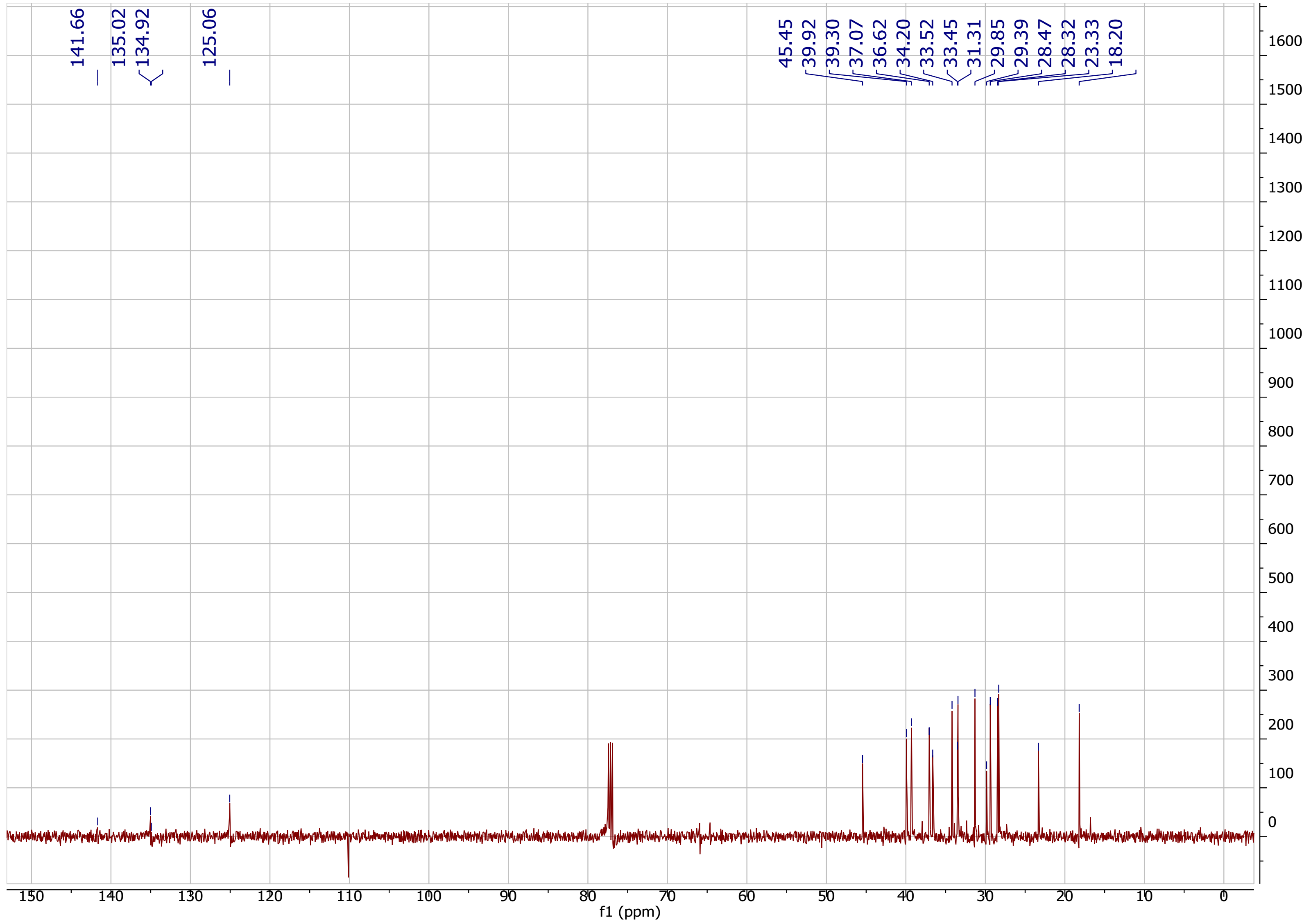


***Supplementary Figure S12.*** $^{13}C$ NMR spectrum (125 MHz) of compound **8** in $CDCl_3$.

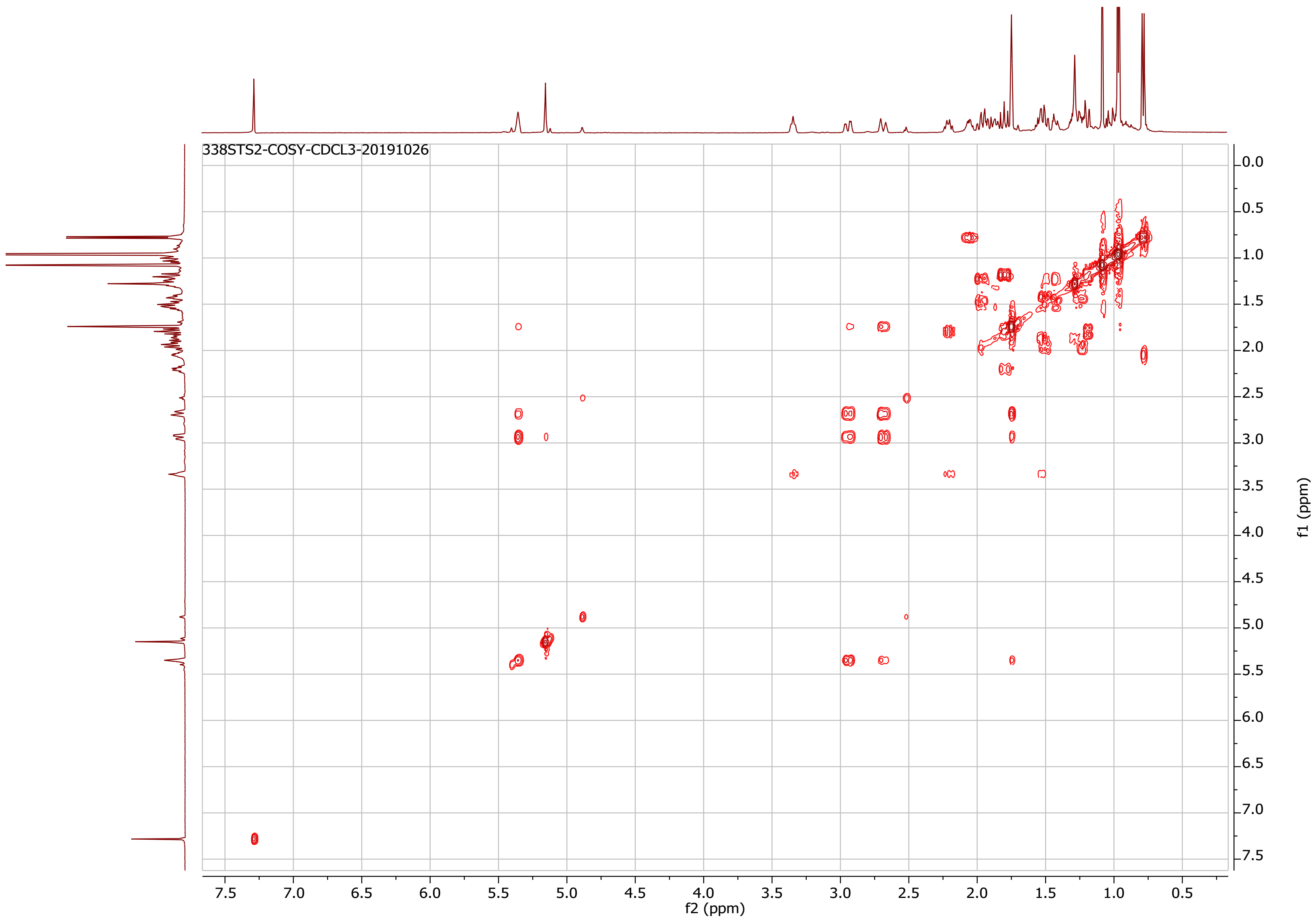


***Supplementary Figure S13.*** [1]H-[1]H COSY spectrum of compound **8** in $CDCl_3$.

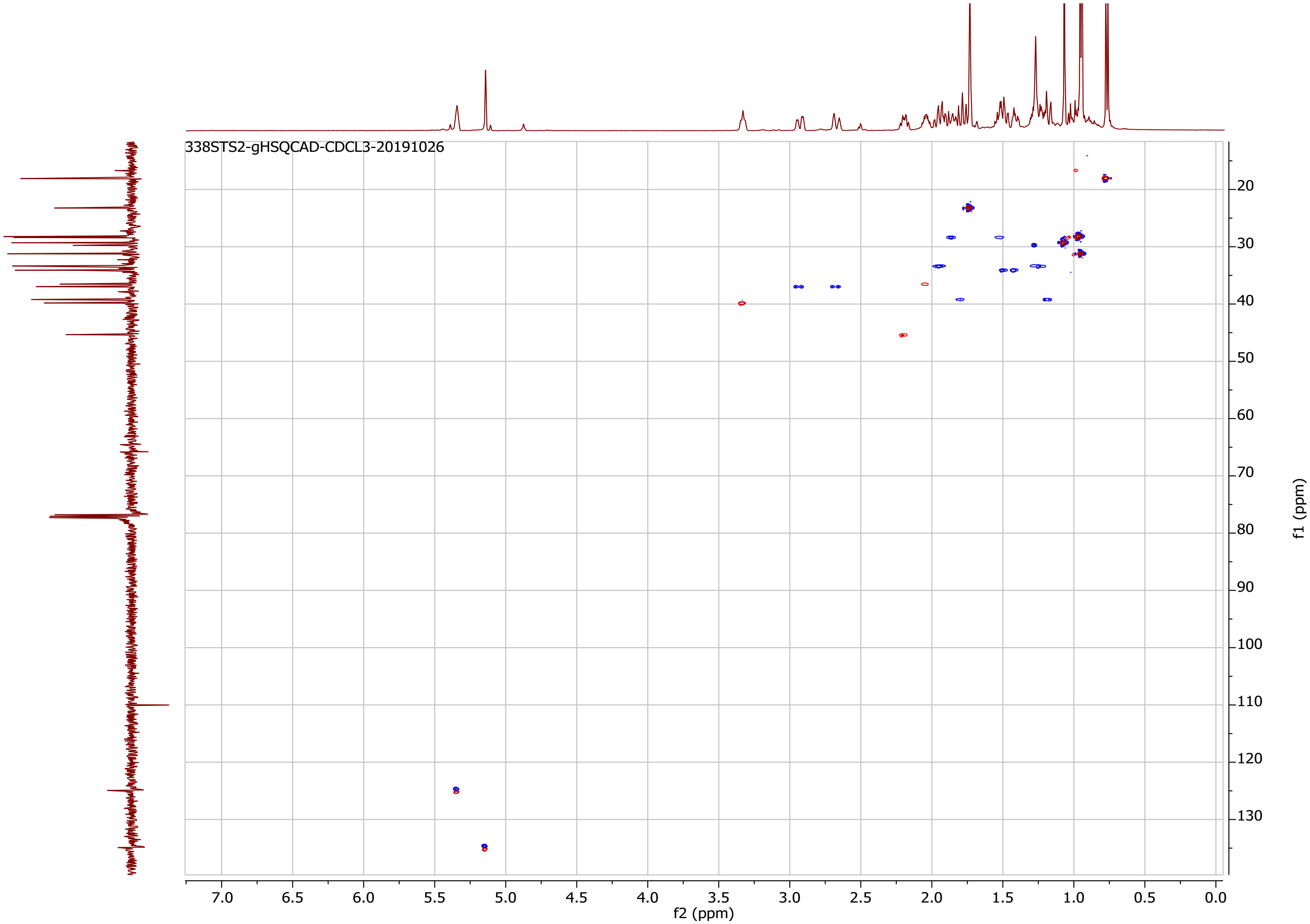


***Supplementary Figure S14**.* HSQCAD spectrum of compound **8** in $CDCl_3$.

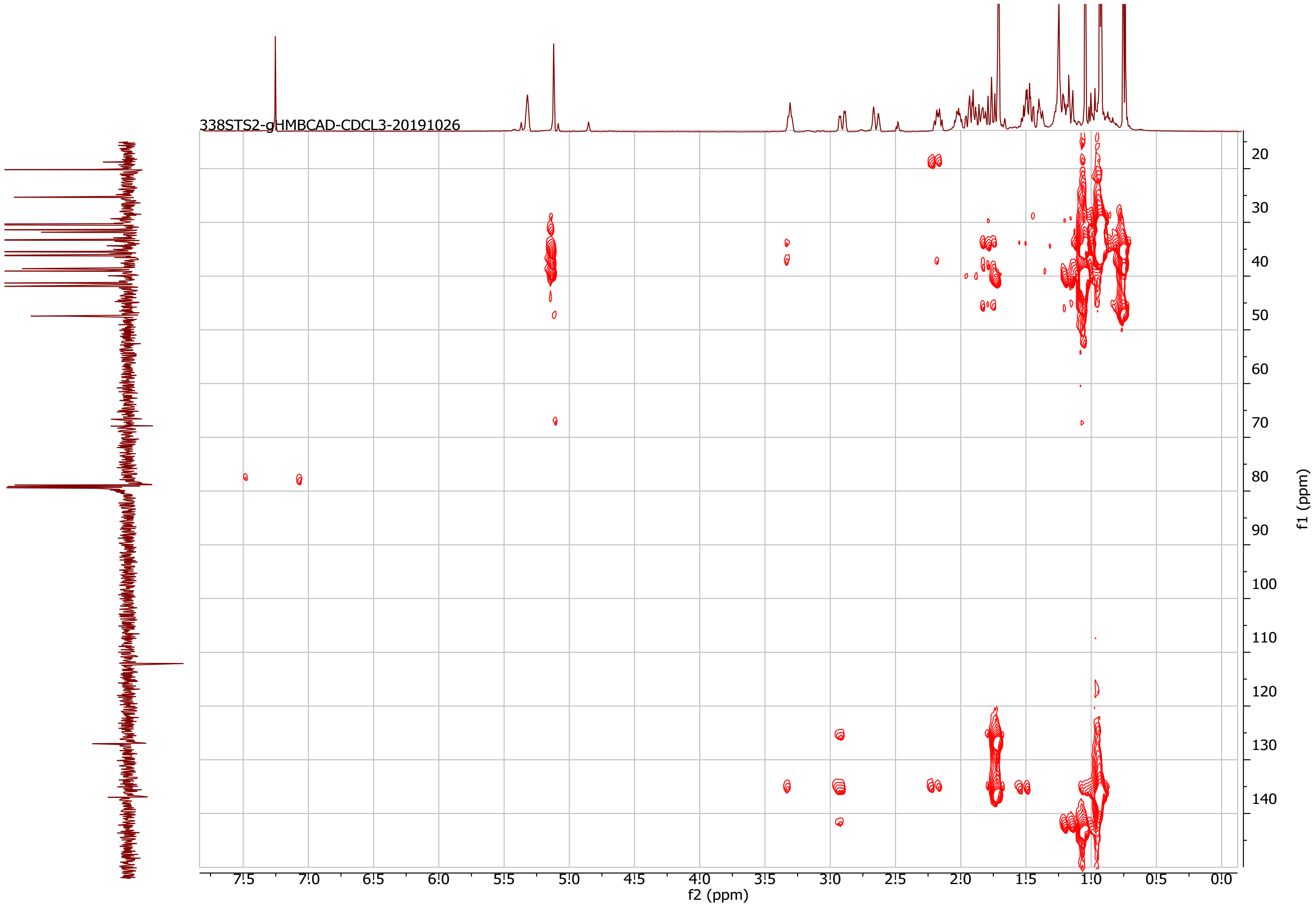


***Supplementary Figure S15.*** HMBCAD spectrum of compound **8** in $CDCl_3$.

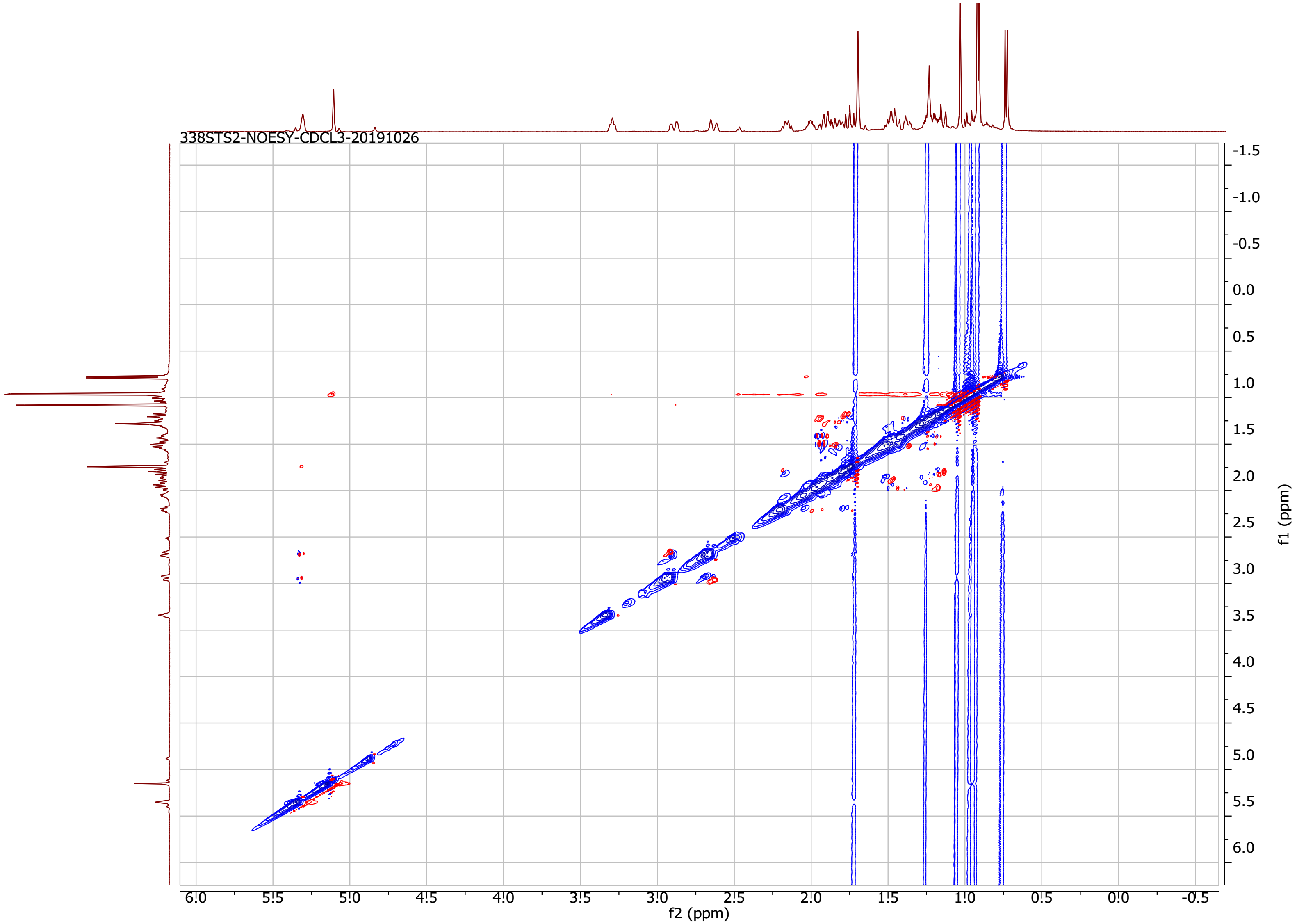


***Supplementary Figure S16***. NOESY spectrum of compound **8** in $CDCl_3$.

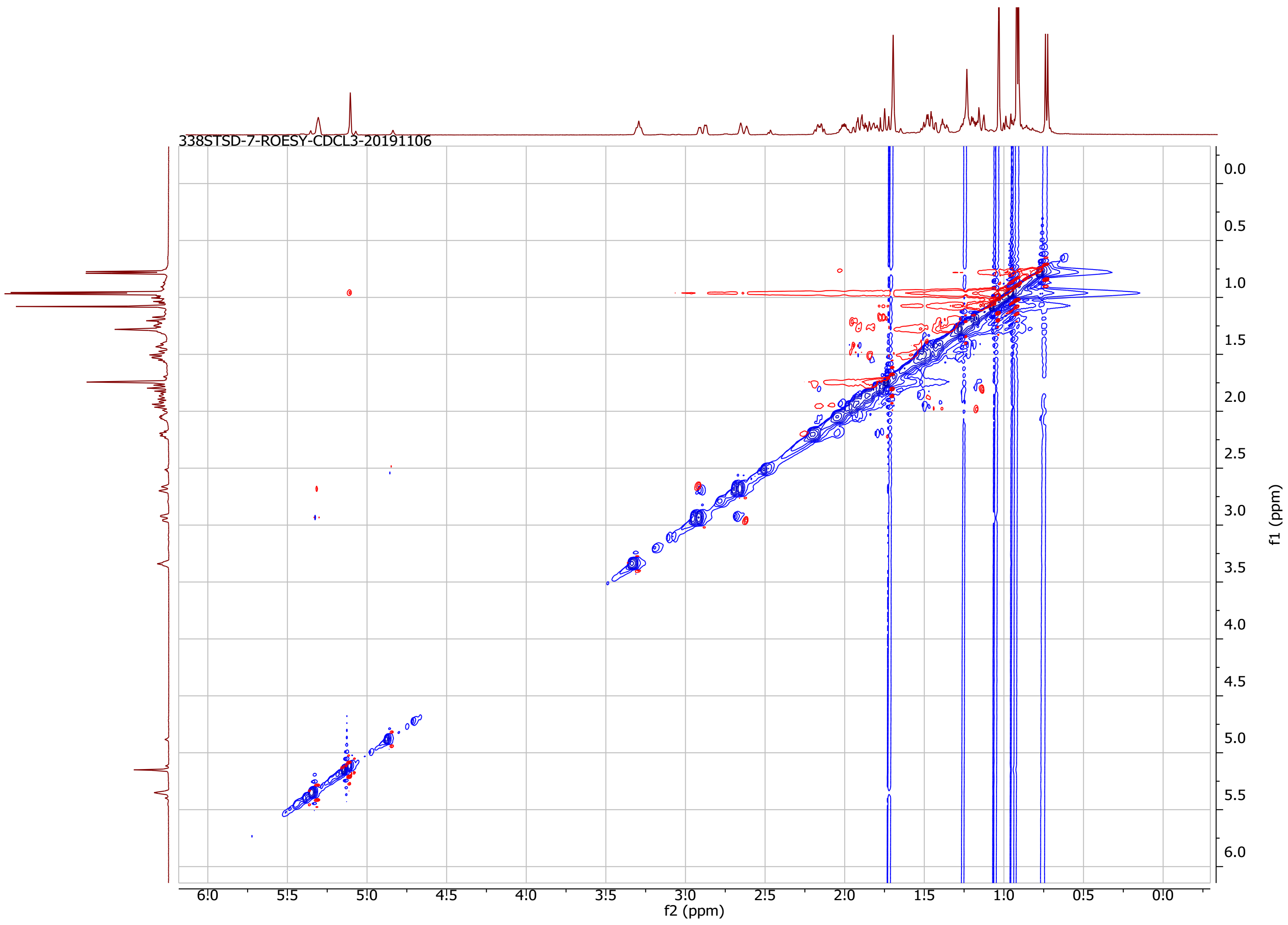


***Supplementary Figure S17***. ROESY spectrum of compound **8** in CDCl3*.*

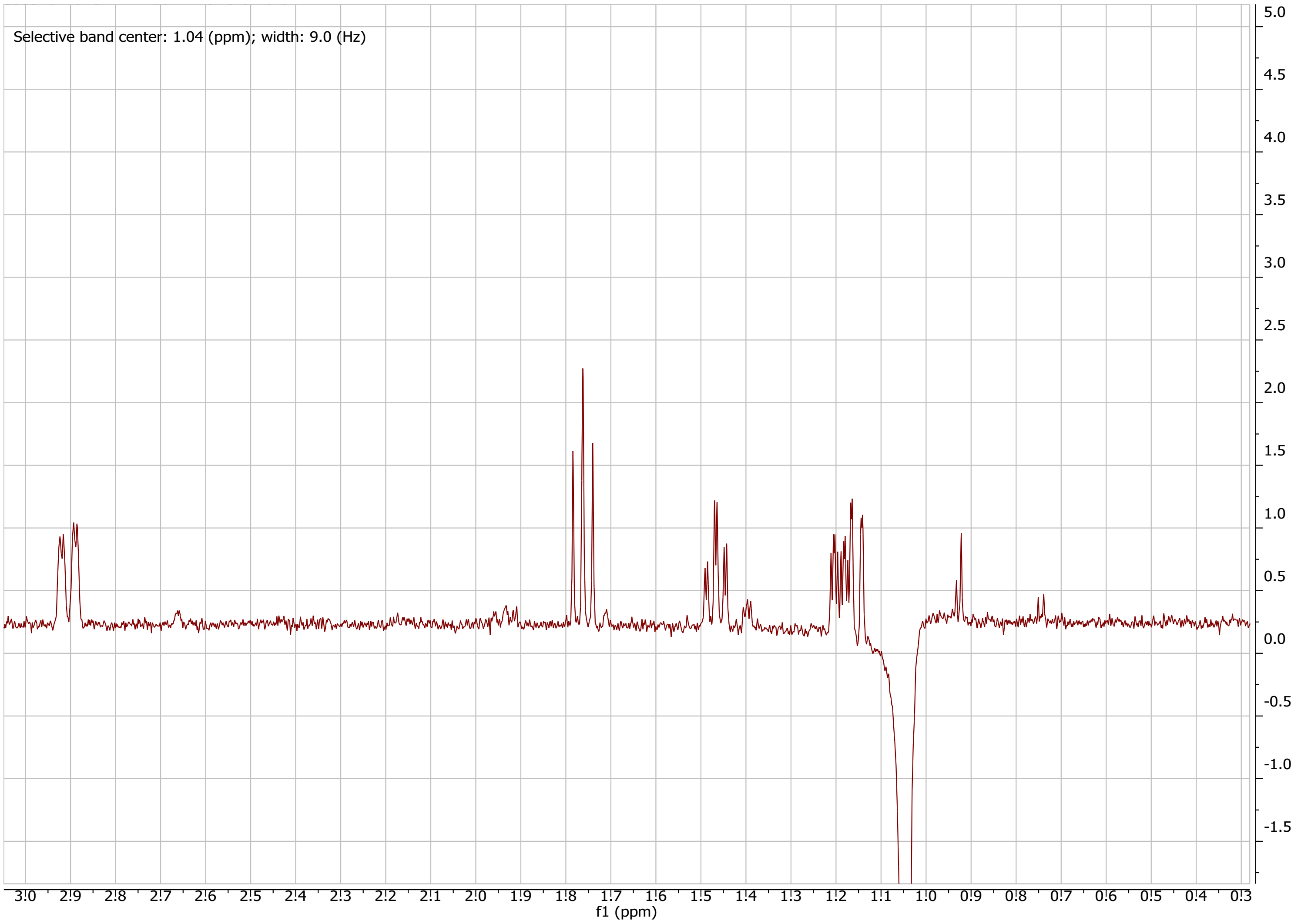


***Supplementary Figure S18.***1D NOESY spectrum of compound **8** in $CDCl_3$ (irradiation at 1.04 ppm).

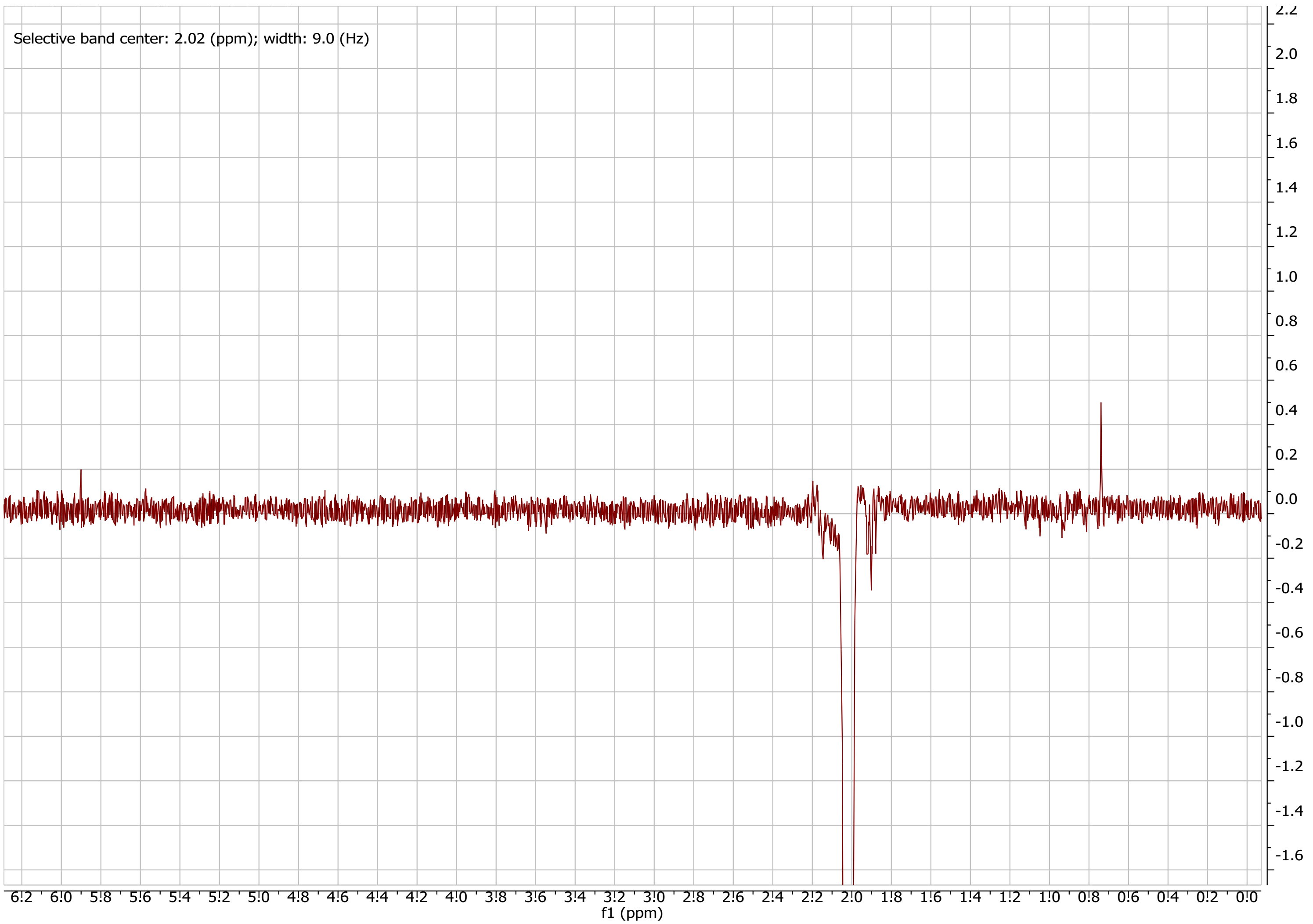


***Supplementary Figure S19.*** 1D NOESY spectrum of compound **8** in $CDCl_3$ (irradiation at 2.02 ppm).

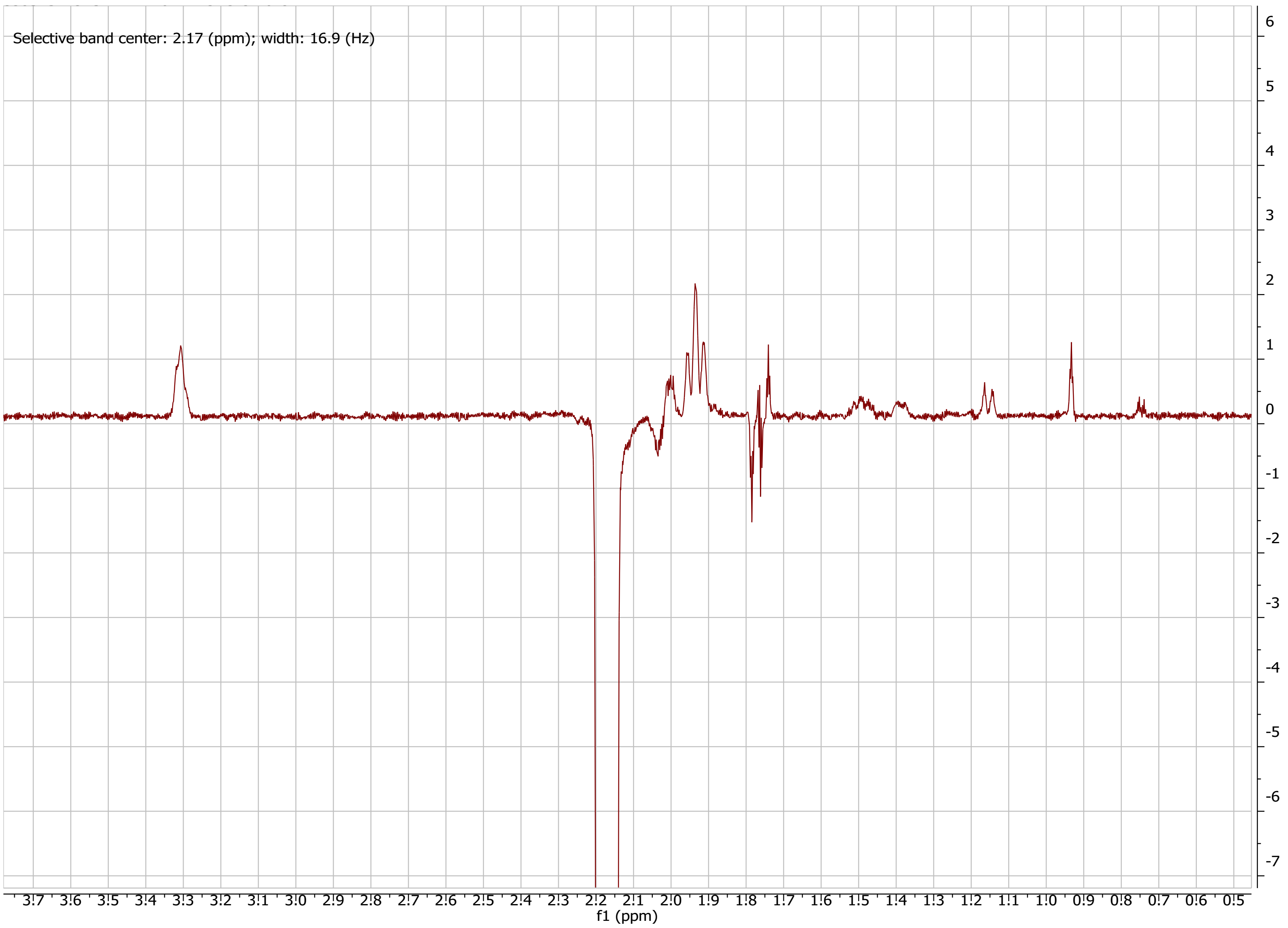


***Supplementary Figure S20.*** 1D NOESY spectrum of compound **8** in $CDCl_3$ (irradiation at 2.17 ppm).

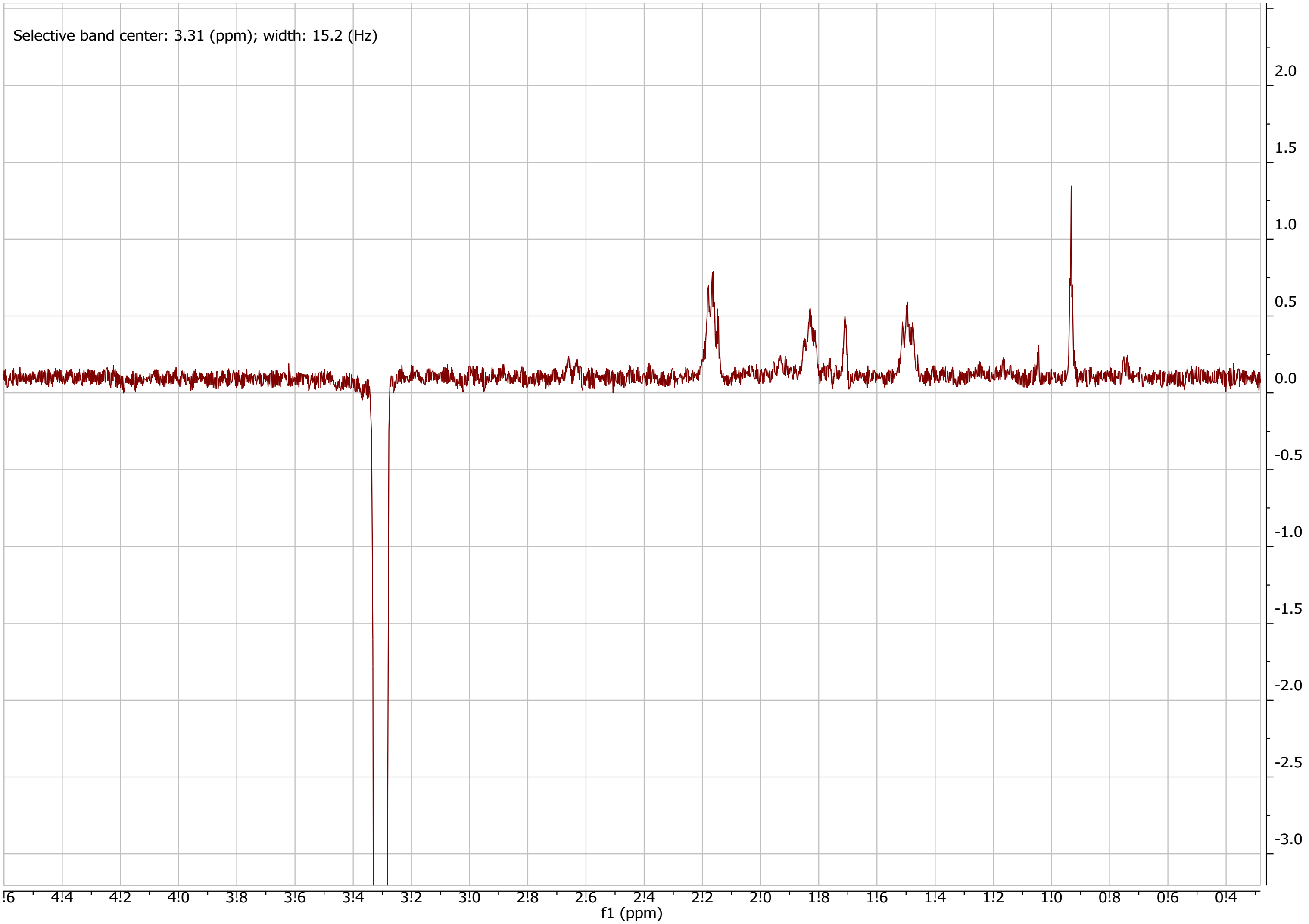


***Supplementary Figure S21***. 1D NOESY spectrum of compound **8** in $CDCl_3$ (irradiation at 3.31 ppm).

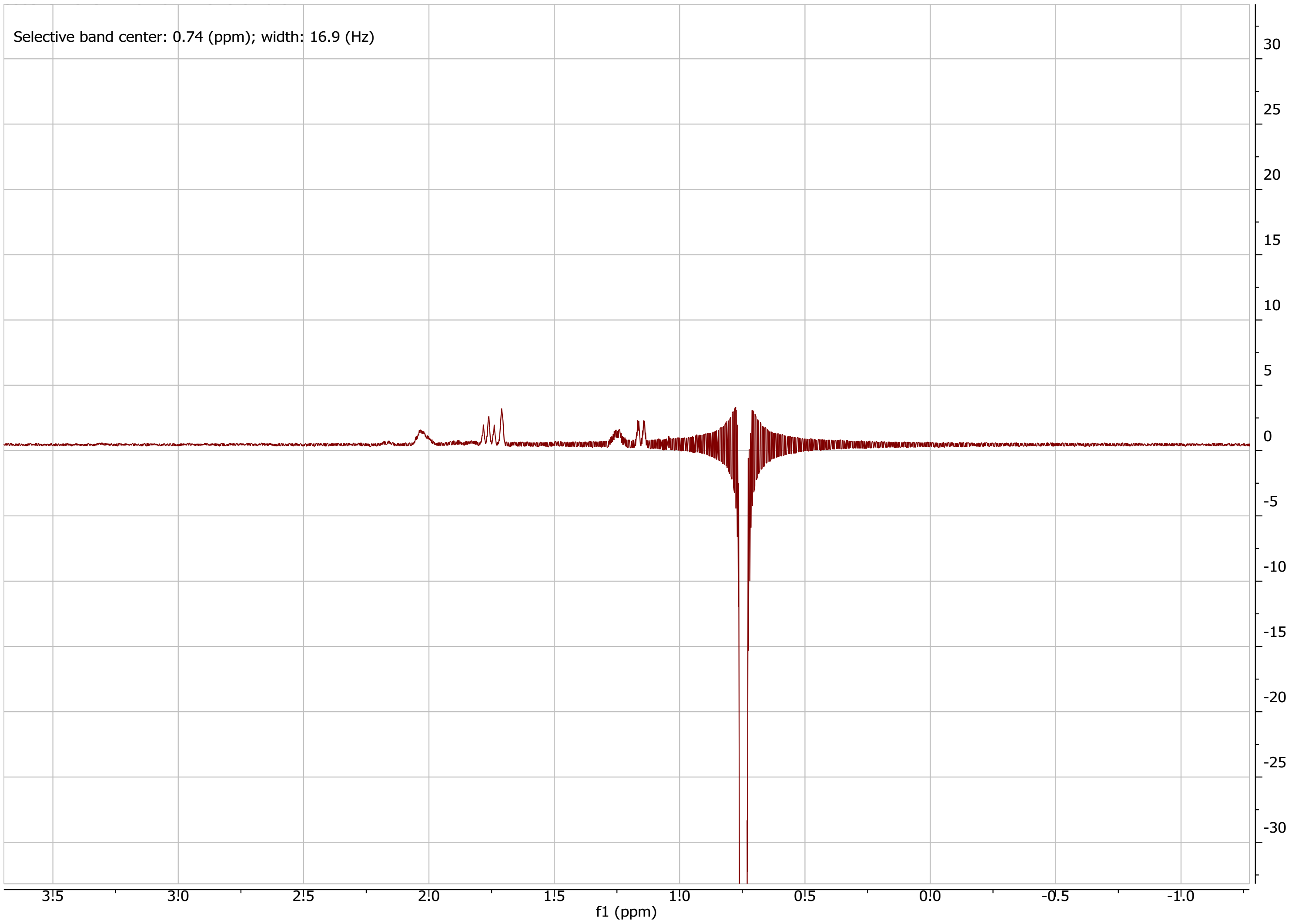


***Supplementary Figure S22***. 1D NOESY spectrum of compound **8** in $CDCl_3$ (irradiation at 0.74 ppm).

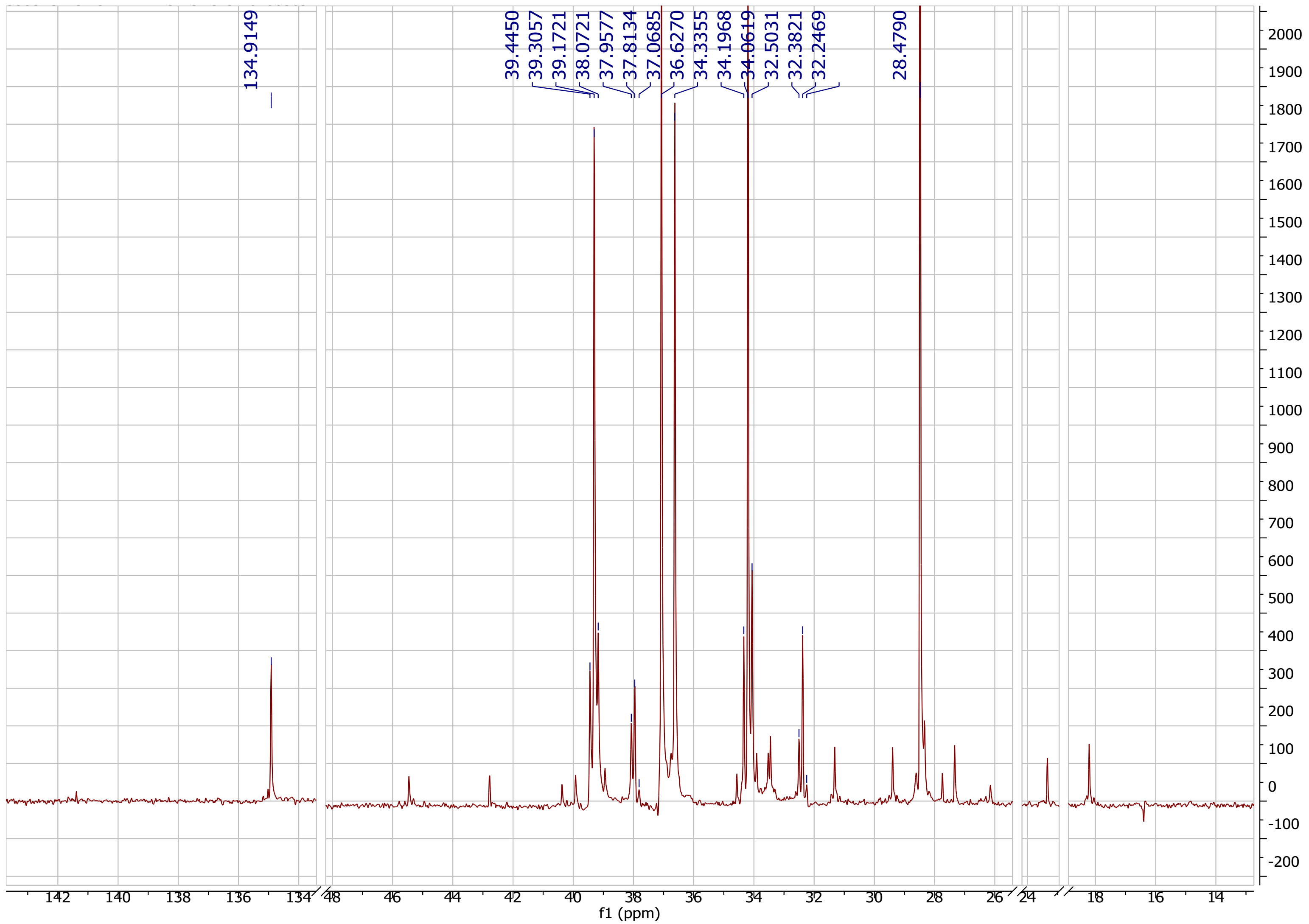


***Supplementary Figure S23.*** $^{13}$C NMR spectrum (125 MHz) of [1-$^{13}$C]acetate-enriched **8** in $CDCl_3$.

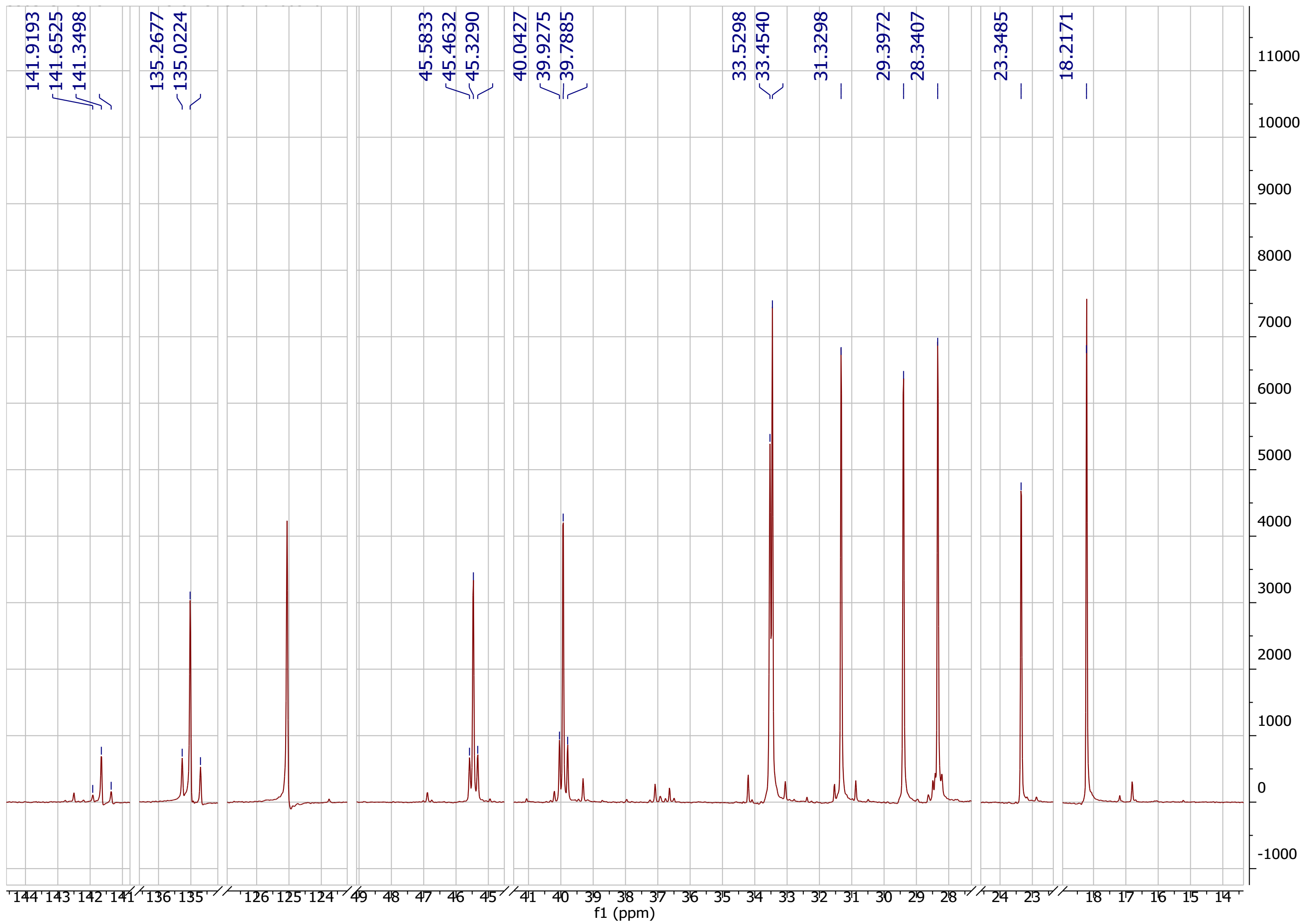


***Supplementary Figure S24.*** $^{13}$C NMR spectrum (125 MHz) of [2-$^{13}$C]acetate-enriched **8** in $CDCl_3$.

***Supplementary Figure S25***. $^{13}$C NMR spectrum (125 MHz) of [1,2-$^{13}C_2$]acetate-enriched **8** in $CDCl_3$. A) Full spectrum to compare resonance intensities; B) Zoomed in areas of the $^{13}$C NMR spectrum to highlight the homonuclear $^{13}$C–$^{13}$C scalar coupling, as well as resonances that are singly enriched. All spectra are shown in ppm with coupling constants shown in Hz.

A)

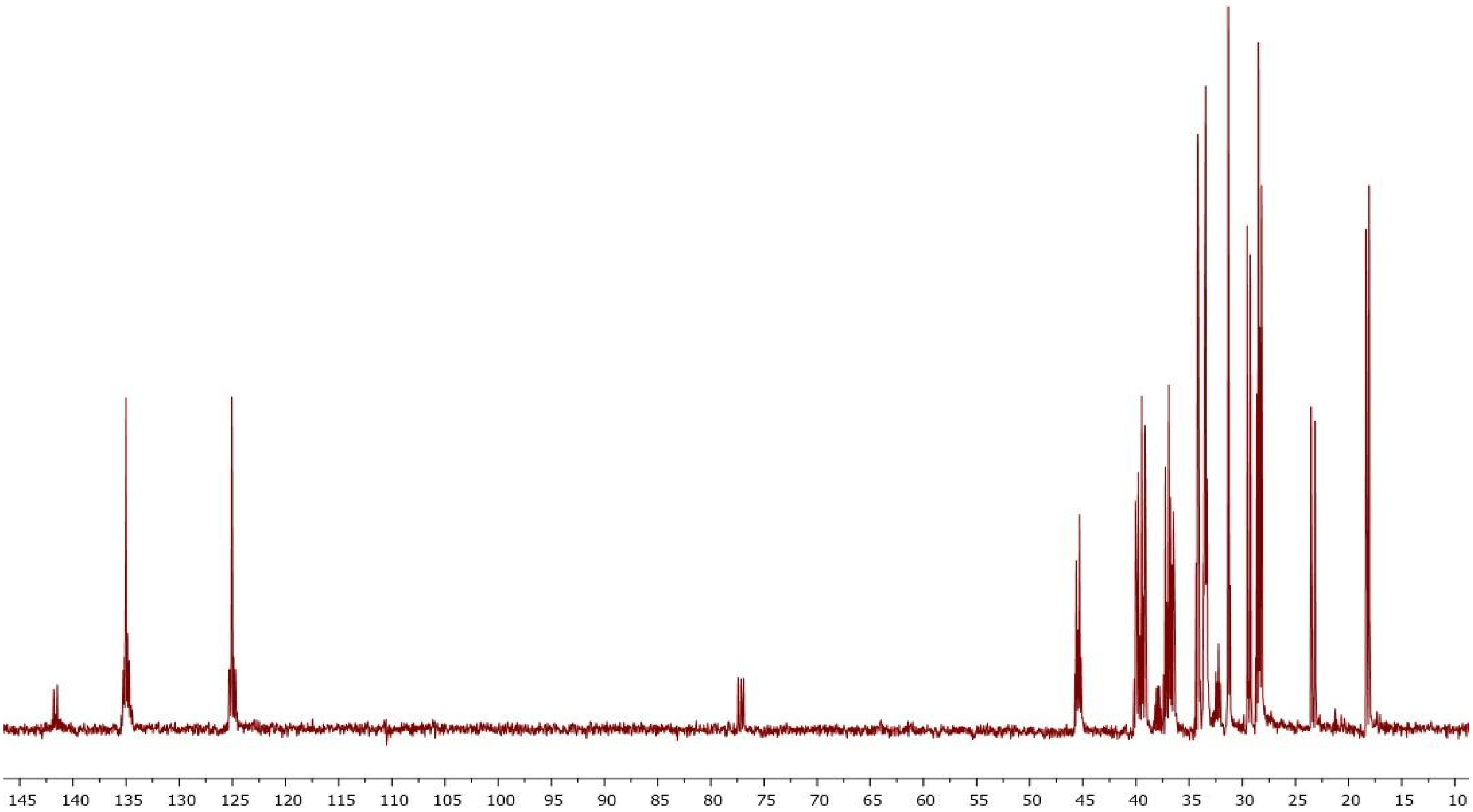

B)

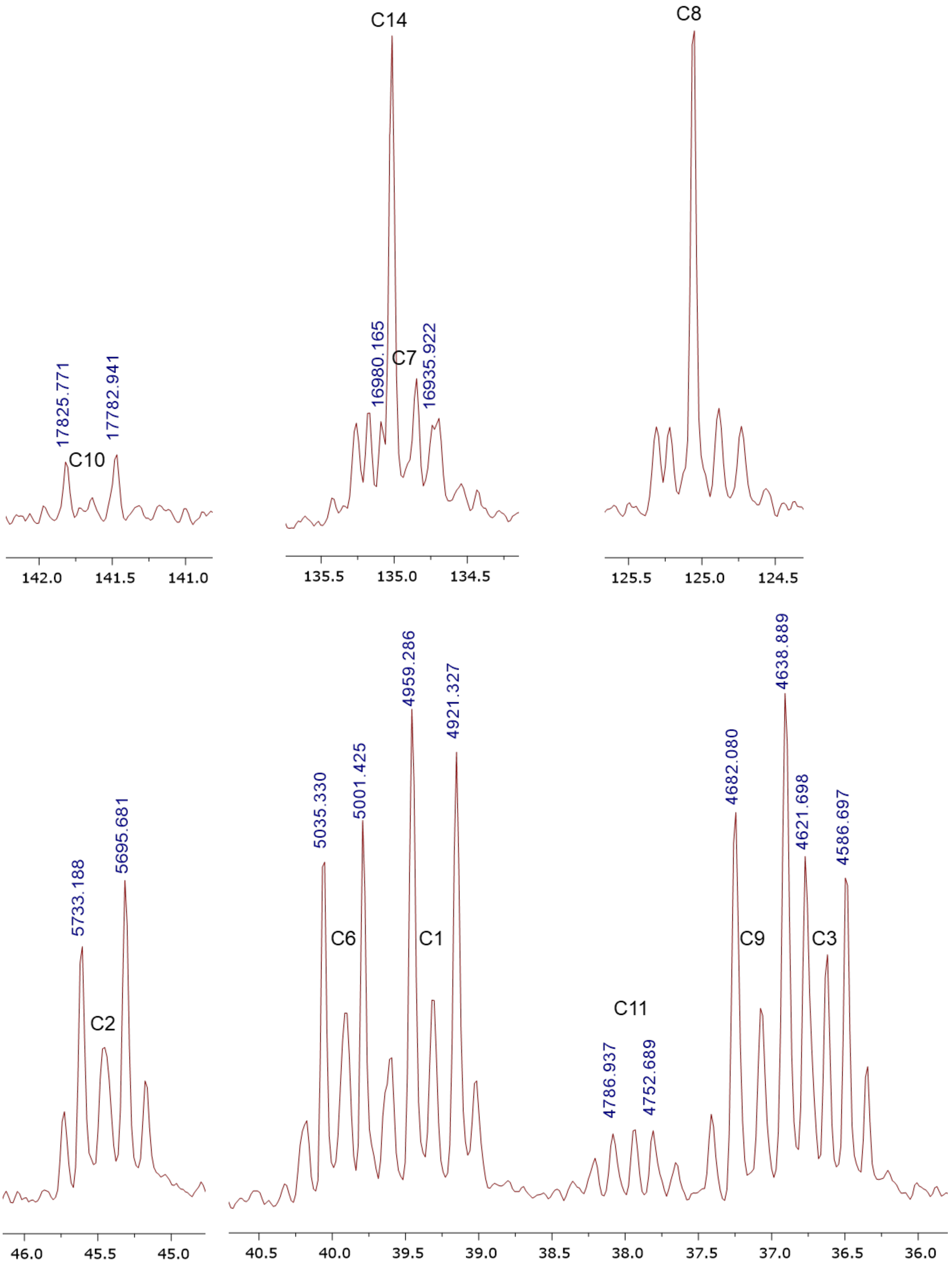

C14
C8
17825.771
17782.941
C10
16980.165
C7
16935.922
142.0
141.5
141.0
135.5
135.0
134.5
125.5
125.0
124.5
4959.286
4921.327
4638.889
4682.080
4621.698
4586.697
5035.330
5001.425
5733.188
5695.681
C6
C1
C9
C3
C2
C11
4786.937
4752.689
46.0
45.5
45.0
40.5
40.0
39.5
39.0
38.5
38.0
37.5
37.0
36.5
36.0

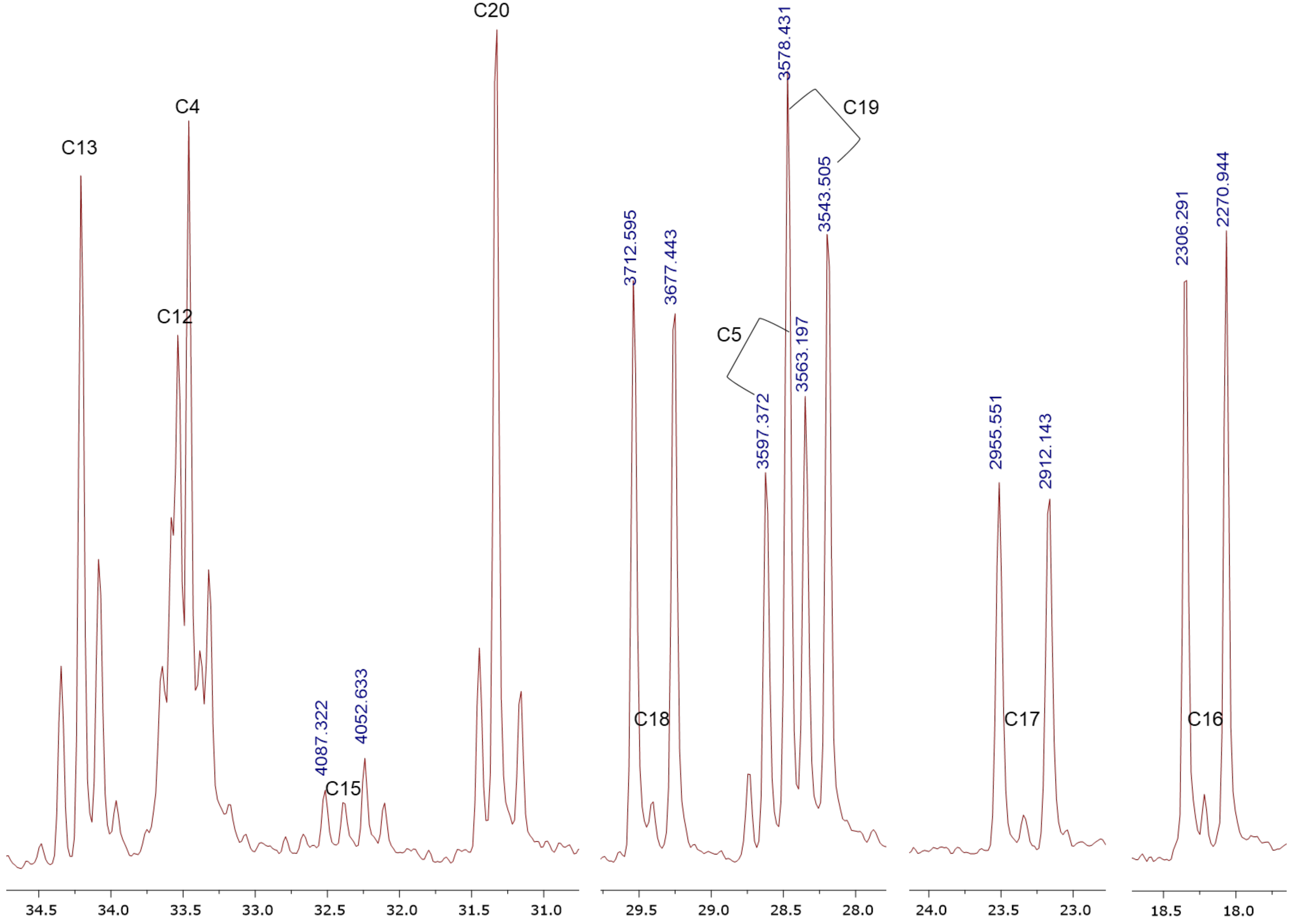
C13
C4
C12
C20
C15
4087.322
4052.633
3712.595
3677.443
C18
3578.431
C19
3543.505
C5
3563.197
3597.372
2955.551
2912.143
C17
2306.291
2270.944
C16
34.5
34.0
33.5
33.0
32.5
32.0
31.5
31.0
29.5
29.0
28.5
28.0
24.0
23.5
23.0
18.5
18.0
